# Tumorigenesis driven by the BRAF^V600E^ oncoprotein requires secondary mutations that overcome its feedback inhibition of migration and invasion

**DOI:** 10.1101/2023.11.21.568071

**Authors:** Sunyana Gadal, Jacob A. Boyer, Simon F. Roy, Noah A. Outmezguine, Malvika Sharma, Hongyan Li, Ning Fan, Eric Chan, Yevgeniy Romin, Afsar Barlas, Qing Chang, Priya Pancholi, Neilawattie. Merna Timaul, Michael Overholtzer, Rona Yaeger, Katia Manova-Todorova, Elisa de Stanchina, Marcus Bosenberg, Neal Rosen

## Abstract

*BRAF^V600E^* mutation occurs in 46% of melanomas and drives high levels of ERK activity and ERK-dependent proliferation. However, *BRAF^V600E^* is insufficient to drive melanoma in GEMM models, and 82% of human benign nevi harbor *BRAF^V600E^* mutations. We show here that BRAF^V600E^ inhibits mesenchymal migration by causing feedback inhibition of RAC1 activity. ERK pathway inhibition induces RAC1 activation and restores migration and invasion. In cells with *BRAF^V600E^*, mutant RAC1, overexpression of PREX1, PREX2, or PTEN inactivation restore RAC1 activity and cell motility. Together, these lesions occur in 48% of BRAF^V600E^ melanomas. Thus, although BRAF^V600E^ activation of ERK deregulates cell proliferation, it prevents full malignant transformation by causing feedback inhibition of cell migration. Secondary mutations are, therefore, required for tumorigenesis. One mechanism underlying tumor evolution may be the selection of lesions that rescue the deleterious effects of oncogenic drivers.

**Statement of significance:** BRAF^V600E^ activation of ERK causes feedback inhibition of cell migration and invasion and thus blocks tumorigenesis. Secondary genetic lesions are required to rescue these processes and enable tumor development. Thus, oncogenic feedback can shape the details of tumor progression and, in doing so, selects for new mutations that may be therapeutic targets.

## Introduction

Mutations that activate ERK signaling, especially those in KRAS, NRAS, and BRAF, are drivers of a substantial fraction of human cancers. The most prevalent BRAF mutants, V600 alleles, have extremely high kinase activity [1] and ERK transcriptional output with concomitant marked ERK-dependent feedback inhibition of upstream signaling and RAS activation [2]. BRAF mutants are common in melanomas, thyroid and colorectal carcinomas, and other types of tumors. Proliferation of these tumors is sensitive to ERK inhibition by MEK or RAF inhibitors and genetic ablation of *BRAF^V600E^* allele [3–5]. This is consistent with the substantial clinical benefit obtained with BRAF inhibitors in BRAF^V600E^ mutant melanomas [3, 6–8].

However, expression of BRAF^V600E^ in melanocytes is not sufficient for the development of melanomas [9–11]. When BRAF^V600E^ was expressed in melanocytes in genetically engineered murine models (GEMMs), it caused epithelial hyperplasia, hyperpigmentation of the skin, and nevi formation but not malignant transformation [10]. In humans, *BRAF^V600E^* mutation is present in 82% of benign nevi, which are pigmented lesions that rarely progress to become malignant melanomas [12]. The Bosenberg and the McMahon laboratories have demonstrated that melanomas do not develop in mice in which either BRAF^V600E^ is expressed, or PTEN is deleted in melanocytes, but BRAF^V600E^ PTEN^null^ mice develop rapidly growing disseminated melanomas [10, 11]. We sought to understand why *BRAF^V600E^* mutation was not sufficient for the development of melanomas, whereas PTEN loss and BRAF^V600E^ together was.

Here, we show that expression of BRAF^V600E^ inhibits cell migration and that pharmacologic inhibition of ERK signaling accelerates the migration and invasion of BRAF^V600E^ and mutant RAS-driven tumors. The suppression of migration and invasion by mutant BRAF is due to ERK-dependent feedback inhibition of RTK signaling, which causes inhibition of RAC1 activation. Thus, feedback inhibition of a key cellular function by the oncoprotein blocks tumorigenesis. In melanoma, *PTEN* loss, activating *RAC1* mutations, or *PREX1/2* amplification co-occur with *BRAF^V600E^* and serve to enhance migration by rescuing RAC1 activation.

## Results

### Cell Migration is inhibited by ERK activation in cells with mutant BRAF

In GEMMs in which BRAF^V600E^ is expressed in melanocytes or lung cells, malignant tumors do not develop [9, 10]. BRAF^V600E^ expression causes cell proliferation followed by growth arrest. In GEMMs, melanocytes with *Braf^V600E^ Cdkn2a^−/−^* mutation formed spatially restricted lesions resembling human melanocytic nevi, with no evidence of melanoma formation. Despite a long latency period, BRAF^V600E^-driven tumors had banal cytological and architectural features and did not infiltrate into lymph nodes or metastasize [10]. In contrast, in GEMMs in which BRAF^V600E^ expression and PTEN, Cdkn2a loss were engineered in melanocytes, invasive melanomas developed quickly and had histopathological features of melanoma, including marked cytological atypia, prominent nucleoli, and atypical mitotic figures. These malignant cells accumulated in the dermis and invaded the subcutis, regional lymph nodes, and formed metastatic nodules in the lung [10]. It was clear that loss of PTEN synergized with hyperactivated RAF to cause tumors. However, the mechanism for the synergy was unknown. In particular, there was no evidence of an increased growth rate of the BRAF^V600E^ PTEN^null^ cells [13]. Our data is consistent with this finding. In fact, the growth rate in tissue culture of melanocytes with BRAF^V600E^ PTEN^null^ appeared to be slower than that of cells with BRAF^V600E^ alone **(Figure S1A)**.

Given these findings, we asked whether BRAF^V600E^ PTEN^null^ cells migrated faster than those with BRAF^V600E^ alone. For these experiments, we used the Yale University Mouse Melanoma (YUMM) cell lines generated from mouse models [13] with genetically defined backgrounds that express BRAF^V600E^ alone or in combination with other alterations that can co-occur with BRAF^V600E^ in human melanomas. To assess the migration and invasion of cell lines in vitro, we used the xCELLigence Real-Time Cell Analyzer (RTCA). The RTCA instrument records cell migration and invasion in real-time and generates cell index curves over time in which an increase in cell index represents the increase in cell migration through a porous membrane of 8μM pore size (migration index) or an increase in cell invasion through the extracellular matrix (ECM) coated membrane (invasion index) [14]. In all migration and invasion assays, regular growth medium was used in the upper and lower chambers of RTCA or transwell assay plates (no gradient).

Cell migration and invasion were assessed in the five YUMM lines used in **S1A.** The *Braf^V600E^ Pten^wildtype^ Cdkn2a^−/−^* model YUMM 3.3 was unable to migrate over 16 hours or invade the ECM by 24 hours (**Figures 1A (i), S1B, S1C)**. Another *Braf^V600E^ Pten^wildtype^ Cdkn2a^−/−^* model, YUMM 3.2, was unable to migrate or invade the ECM for the first 8 hours. After a lag of 8 hours, 3.2 starts to migrate and invade. In contrast, two YUMM lines (1.3, 1.1), with *Braf^V600E^* and homozygous deletion of *Pten,* migrated and invaded through the ECM rapidly. Migration indices of 1.3, 1.1, and 3.2 were 775%, 428%, and 83% higher than that of YUMM 3.3 at 8 hours (**Figure 1A (i)**). Although 3.2 migrated and invaded faster than 3.3, it was still slower than 1.1 and 1.3. Of note, 3.2 expresses 51% less PTEN protein than 3.3 (**Figure 1A (ii)**). YUMM 4.1 (homozygous deletion of *Pten* and wildtype *Braf)* migrated and invaded the ECM even faster than and 1.3, and at 8 hours, its migration index was 1178% higher than that of YUMM 3.3. Transwell assays through ECM-coated upper chambers corroborated the findings obtained with the RTCA invasion assay **(Figure S1C)**. These data suggest that either melanocytes do not migrate well enough to support the transformed phenotype, and the loss of PTEN enhances migration in cells with mutant or wildtype (WT) BRAF, or that activated BRAF suppresses melanocyte mobility and that this suppression is rescued by PTEN loss.

**Figure 1.**
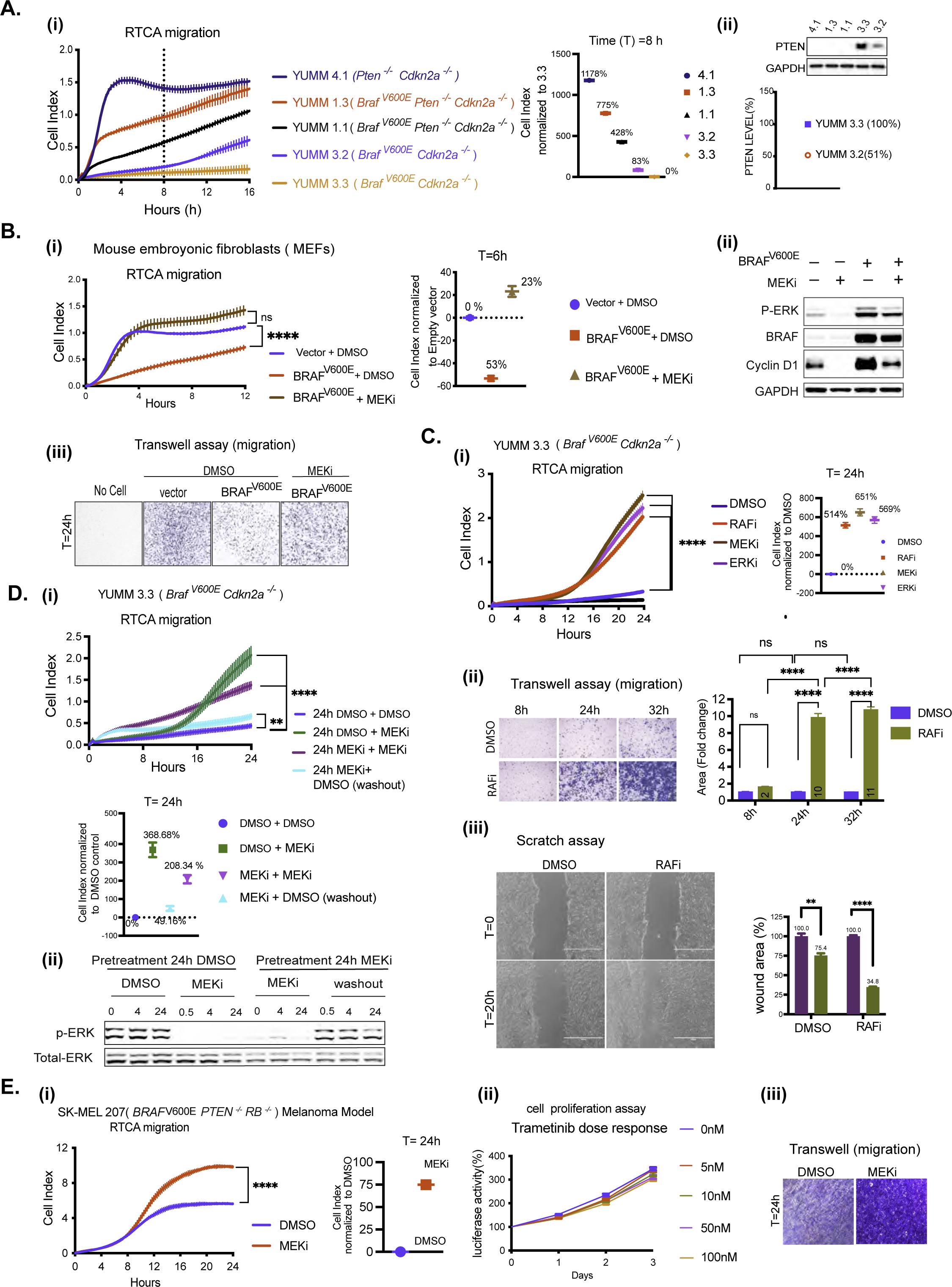
BRAF^V600E^-driven ERK signaling inhibits cell migration in vitro. **A. i)** Migration curves of YUMM 4.1, 1.3, 1.1, 3.3, and 3.2 generated by RTCA. Cell index at 8 hours(h) was normalized to YUMM 3.3. ii) Whole cell lysate (WCL) was analyzed by immunoblotting (IB) to determine PTEN levels. Bands were quantified by densitometry, and the PTEN level was normalized to YUMM 3.3. **B. i-iii)** MEFs were transfected with empty vector (Vector) or *BRAF*^V600E^ plasmid and 36h after transfection, seeded for RTCA, transwell assay and IB. **i)** Migration with Dimethyl sulfoxide (DMSO) or 10nM Trametinib (MEKi). P-values are based on ordinary one-way ANOVA analysis. Cell index at 6h was normalized to vector. **ii)** ERK signaling was assayed by IB. **iii)** Transwell assay was performed with DMSO or 10nM Trametinib and visualized by crystal violet staining (CV) at 24h (n=3). **C. i)** Migration of YUMM 3.3 was assessed with DMSO, 1uM Vemurafenib (RAFi), 10nM Trametinib (MEKi), or 500nM SCH 772984 (ERKi) treatment. P-values are based on ordinary one-way ANOVA analysis. Cell index at 24h was normalized to DMSO control. **ii)** Transwell assay 1uM Vemurafenib was performed as in 1B. Bar graphs depict area with positive CV stain, Error bars mean ± SEM (n=3). P-values are based on two-way ANOVA analysis. **iii)** YUMM 3.3 cells were plated with DMSO or 1uM Vemurafenib 24 hours before wounding. Bright-field image of the scratch was taken at T=0h (within 30 minutes of wounding) and T=20h. The bar graph depicts the open wound area. Error bars mean ± SE (n=3). P-values are based on multiple unpaired t-tests. DMSO control. **ii)** Transwell assay 1uM Vemurafenib was performed as in 1B. Bar graphs depict area with positive CV stain, Error bars mean ± SEM (n=3). P-values are based on two-way ANOVA analysis. **iii)** YUMM 3.3 cells were plated with DMSO or 1uM Vemurafenib 24 hours before wounding. Bright-field image of the scratch was taken at T=0h (within 30 minutes of wounding) and T=20h. The bar graph depicts the open wound area. Error bars mean ± SE (n=3). P-values are based on multiple unpaired t-tests. **D. i)**YUMM 3.3 cells were pre-treated with DMSO or 10nM Trametinib(24h). Pre-treated cells were reseeded with either DMSO or 10nM Trametinib for RTCA, and cell index recorded. To wash out the drug, Trametinib pretreated cells were washed with phosphate buffer solution (PBS) three times before seeding in RTCA chambers with DMSO. P-values are based on ordinary one-way ANOVA analysis. Cell index at 24h was normalized to control (24h DMSO + DMSO). **ii)** Cell pellets were collected at indicated times, and ERK signaling was analyzed by IB. **E. i)** Migration of SK-MEL207 with DMSO or 10nM Trametinib. Cell index at 24h was normalized to DMSO control. P-values are based on two-tailed unpaired Student’s t-test **ii)** Cells were seeded in 96 well plates and treated with DMSO or Trametinib. At indicated times, cell growth was measured using CellTiter-Glo Cell Viability Assay. Error bars mean ± SD (n=6). iii) Transwell assay, was performed with 10nM Trametinib(24h) treatment and visualized by CV(n=3). Migration curves with vertical error bars represent mean ± SEM (n=3). *P*<0.0001 is shown as ****, P<0.001 as ***, P< 0.01 as **, and *P*>0.05 as ns. Also, see SF1.

To address this question, BRAF^V600E^ was expressed in mouse embryonic fibroblasts (MEFs) that migrate rapidly (**Figure 1B (i))**. Expression of BRAF^V600E^ in these cells increased ERK phosphorylation and reduced their mobility by 53%. Inhibition of migration was proportional to levels of BRAF^V600E^ expression and ERK phosphorylation **(Figure S1D)**. Trametinib, a selective inhibitor of MEK, inhibited ERK signaling and restored the migration of BRAF^V600E^ expressing MEFs (**Figures 1B (i), (ii), (iii))**. These data suggest that activation of ERK by BRAF^V600E^ suppresses the migration of melanocytes.

Consistent with this hypothesis, inhibition of ERK signaling with RAF, MEK, or ERK inhibitor stimulates the migration of the YUMM 3.3 (*Braf^V600E/+^Cdkn2a^−/−^*) cells (**Figure 1C (i))**. Migration is induced 8-12 hours after drug addition. The cell index by 24 hours was increased by approximately 500% over DMSO control by each ERK pathway inhibitor. Similarly, in a transwell assay, 24 hours after treatment, Vemurafenib (RAFi) increased cell migration approximately 9-fold. (**Figure 1C (ii))**. Similarly, this induction was corroborated in a wound-healing assay. After 20 hours of wounding, 34% of the wounded area was left open with Vemurafenib-treated YUMM 3.3 cells, whereas 75% was open with control cells. (**Figure 1C (iii))**. To determine whether the effect of ERK inhibition was reversible, YUMM 3.3 was treated with Trametinib or DMSO. After 24 hours, pretreated cells were seeded for RTCA with fresh DMSO or Trametinib-containing medium (**Figure 1D**). Migration of cells pretreated with DMSO followed by Trametinib was increased by 368% compared to the control (DMSO+DMSO) by 24 hours. In MEK inhibitor pretreated cells replated in Trametinib containing fresh medium, ERK signaling remained inhibited, and migration increased by 208% compared to control. Washing out the drug from MEKi pretreated cells rapidly increased ERK phosphorylation to pretreatment levels (**Figure 1D (ii))**. When these cells were seeded with DMSO-containing media, migration was increased by only 49% compared to the control. Thus, in YUMM 3.3, MEK inhibition enhances migration; this effect is reversed when the drug is washed out and ERK is reactivated.

This phenomenon occurs in multiple cancer cell models with mutant activation of ERK signaling. ERK inhibition enhances migration **(Figure S1I)** in all of the BRAF^V600E^ YUMM and human melanoma cell lines we tested and in the BRAF^V600E^ thyroid cancer cell line B47275 derived from a GEMM model **(Figures S1E, S1F, S1G)**. Even though BRAF^V600E^ PTEN^null^ YUMM and melanoma models **(Figure S1F)** migrate effectively, their migration is increased by MEK inhibition. Thus, in tumor cells with BRAF^V600E^, suppression of migration by ERK is a general phenomenon that is relieved by ERK pathway inhibition. Although loss of PTEN rescues BRAF^V600E^ suppression of migration, ERK pathway inhibition enhances migration further.

Tumors with mutant NRAS or NF1 loss activate the RAS pathway. RAS activation drives both ERK and PI3K signaling in tumors, and PI3K signaling is a well-known inducer of cell motility [15]. We found NRAS mutant and NF1 null melanoma cells migrate well, but MEK inhibition further enhances their migration **(Figure S1H)**. These results suggest that the migration rate is a balance between activating and inhibitory pathways, of which ERK signaling is one of the latter.

The proliferation of BRAF mutant tumors is ERK-dependent [3], and the proliferation of the models studied here is arrested 2-3 days after exposure to the drug. **(Figures S1J, S1K)**. We asked whether induction of migration by ERK pathway inhibition is due to inhibition of proliferation. Inhibition of proliferation of BRAF^V600E^ melanoma cells is preceded by cyclin D/cdk4/6 inhibition and is RB-dependent. Growth of RB-negative BRAF^V600E^ melanomas is insensitive to ERK inhibition by ERK pathway inhibitors [16]. We examined the migration of the RB-negative SK-MEL 207 cell line (*BRAF^V600E/+^ PTEN^−/−^ RB^−/−^*). Consistent with the absence of PTEN, these cells migrated significantly.10nM Trametinib enhanced their migration further (**Figure 1E (i))** but had no effect on cell proliferation at concentrations of up to 100nM (**Figure 1E (ii)).** Similar results were obtained in a transwell migration assay with 10nM Trametinib at 24 hours (**Figure 1E (iii))**. Thus, the induction of migration by ERK inhibition does not depend on the slowing of cell proliferation.

### Inhibition of RAF enhances invasion of BRAF^V600E^ tumors *in vivo*

YUMM 3.3 cells do not appreciably invade through ECM, but invasion was stimulated 8-12 hours after exposure to Vemurafenib (**Figure 2A (i))**. ERK inhibition enhanced invasion of BRAF^V600E^ YUMM models (3.2, 1.1) and human melanoma cell lines (SK-MEL207 and A375) **(Figure S2A)**. As proliferation of SK-MEL207 is unaffected by ERK pathway inhibition (**Figure 1E**), increased invasion is not due to growth inhibition. Results obtained through RTCA were verified by transwell assay. (**Figures 2A(ii), S2A)**.

**Figure 2.**
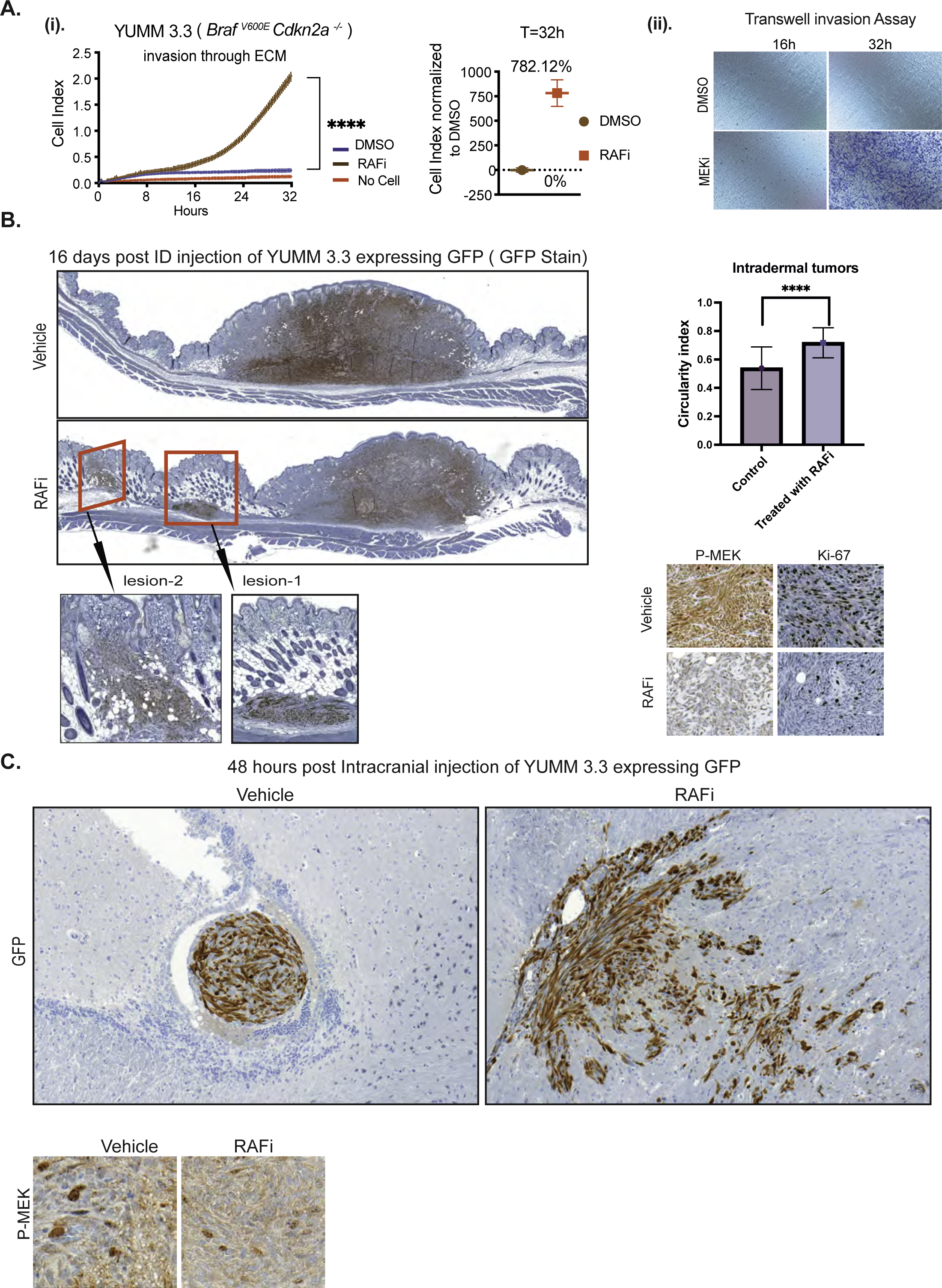
BRAF^V600E^-driven ERK signaling inhibits migration and invasion in vivo. A. **i)** Invasion curve of YUMM 3.3 was generated using ECM-coated CIM plates with DMSO or 1uM Vemurafenib treatment. P-values are based on ordinary one-way ANOVA analysis. The invasion index at 32h was normalized to DMSO control. **ii)**, ECM-coated transwell plates were used, and invasion was assessed followed by CV staining (n=3). **B.** GFP-expressing YUMM 3.3 cells were pre-treated with either DMSO or 1uM RAF inhibitor (PLX 4720) for 24 hours in culture and injected intradermally. Vehicle or 7.5mg/kg PLX 4720 BID was started 24 hours post-injection. Sixteen days post-injection, tumors with surrounding tissue were collected, and histological sections were stained with GFP, Ki67, and phospho-MEK antibodies. The red box shows lateral extensions from the primary larger tumor nodule of YUMM 3.3 cells. The inset displays microsatellitosis at higher magnification. Bar graph depicts the circulatory index. **C.** GFP-expressing YUMM 3.3 cells were pre-treated with DMSO or 1uM Vemurafenib for 24 hours in culture and injected intracranially. Forty-eight hours post-injection, brain was collected, and histopathological sections were stained with GFP and phospho-MEK antibodies. Invasion curves with vertical error bars represent mean ± SEM (n=3). *P*<0.0001 is shown as ****, P<0.001 as ***, P< 0.01 as **, and *P*>0.05 as ns. Also, see SF2.

We asked whether ERK signaling suppresses migration and invasion *in vivo*. YUMM 3.3 cells that express green fluorescence protein (GFP) were injected intradermally in C57BL/6 mice, and twenty-four hours post-injection, either 7.5mg/kg (BID RAF inhibitor PLX 4720 (closely related to vemurafenib but easier to formulate) or vehicle were administered until the day tumors were resected. Local neoplasms developed in mice within seven days of injection **(Figure S2B)**. These masses were comprised of spindle cells in interlacing fascicles. In mice treated with vehicle, YUMM 3.3 formed a single solitary mass (**Figures 2B, S2B)** at the primary site of injection, and there was no evidence for lateral spread of the cells by sixteen days post-injection. By contrast, seven days after injection, 25% of mice treated with PLX 4720 had secondary lateral tumor growths distinctly separated from the larger primary nodule without intervening inflammation or fibrosis (red box **Figure S2B),** reminiscent of the phenomenon of microsatellitosis in human melanoma. Microsatellitosis is a histopathological reporting parameter that portends a poor prognosis in human patients with melanoma [17]. Sixteen days after injection, 70% of animals receiving RAF inhibitor had tumors with microsatellites (range: 1-3 microsatellites per tumor) (red box **Figures 2B, S2B)** the inset shows secondary foci at higher magnification. Microsatellites were located at a mean of 985±860μm from the primary tumor nodule. We subsequently evaluated the circularity/roundness of annotated tumor image objects to estimate whether tumor surface area and perimeter approximated a circle or an irregular shape with an irregular invasive front, as suggested by the circularity index (CI, closer to 1 indicates that the shape approximates a circle). The CI of intradermal melanomas was greater in RAFi-treated cases (**Figure 2B**). This is likely attributable to the phenomenon of microsatellites, which in human melanoma usually manifest as well-circumscribed, round dermal malignant nodules without epidermal connection, akin to in-transit or distant metastases. Ki67 and p-MEK staining were significantly inhibited with RAF inhibitor treatment (**Figure 2B**). Thus, both migration and invasion of YUMM 3.3 were enhanced by ERK inhibition *in vivo*. GFP-expressing YUMM 3.3 cells pretreated with either DMSO or 1uM PLX4720 (24h) were injected intracranially. Forty-eight hours after injection, control cells gave rise to neoplastic cell aggregates of various sizes in the cerebral ventricle that did not invade the parenchyma and remained confined by the thin ependymal cell lining of the ventricular cavity. In contrast, vemurafenib pretreated cells had infiltrated through the ependymal lining and into the brain parenchyma (**Figures 2C, S2C**). They were dispersed within the brain mass, in a pattern of nodules with irregular tumor borders and single cells, at a mean intraparenchymal depth of 660±206 μm from the ependymal cell lining of the ventricle. Similar levels of MEK phosphorylation were detected in cells derived from DMSO and RAF inhibitor pretreated cells; this is not surprising since mice did not receive any further treatment post-injection. (**Figure 2C).** These results illustrate that inhibition of ERK signaling enhances the invasion of BRAF^V600E^ melanocytes both in tissue culture and *in vivo*, inducing *in vivo* microsatellitosis reminiscent of human melanoma and intraparenchymal brain invasion beyond the ependymal lining.

### Cells with BRAF^V600E^ acquire mesenchymal traits after ERK pathway inhibition

Cells can migrate as multicellular sheets, detached clusters, or single cells that employ mesenchymal or amoeboid movement [18–20]. We asked whether induction of migration by ERK pathway inhibition was associated with changes in cell morphology. YUMM 3.3 and YUMM 3.2 cells form multicellular units, with each cell forming junctions with adjacent cells (black arrow, **Figures 3A, S3A)**. Within 24 hours of ERK inhibition, cell-cell junctions were lost, and the multicellular units were replaced by single cells or by a population of loosely scattered elongated cells (red arrow, **Figures 3A, S3A).** These results suggest that ERK inhibition stimulates the mesenchymal mode of motility [18–20]. The mesenchymal transition involves the loss of intercellular junctions, loss of apical-basal polarity, and restructuring of the cytoskeleton [21] with the formation of actin-rich membrane protrusions (lamellipodia) at the cell’s leading edge [22, 23].

**Figure 3:**
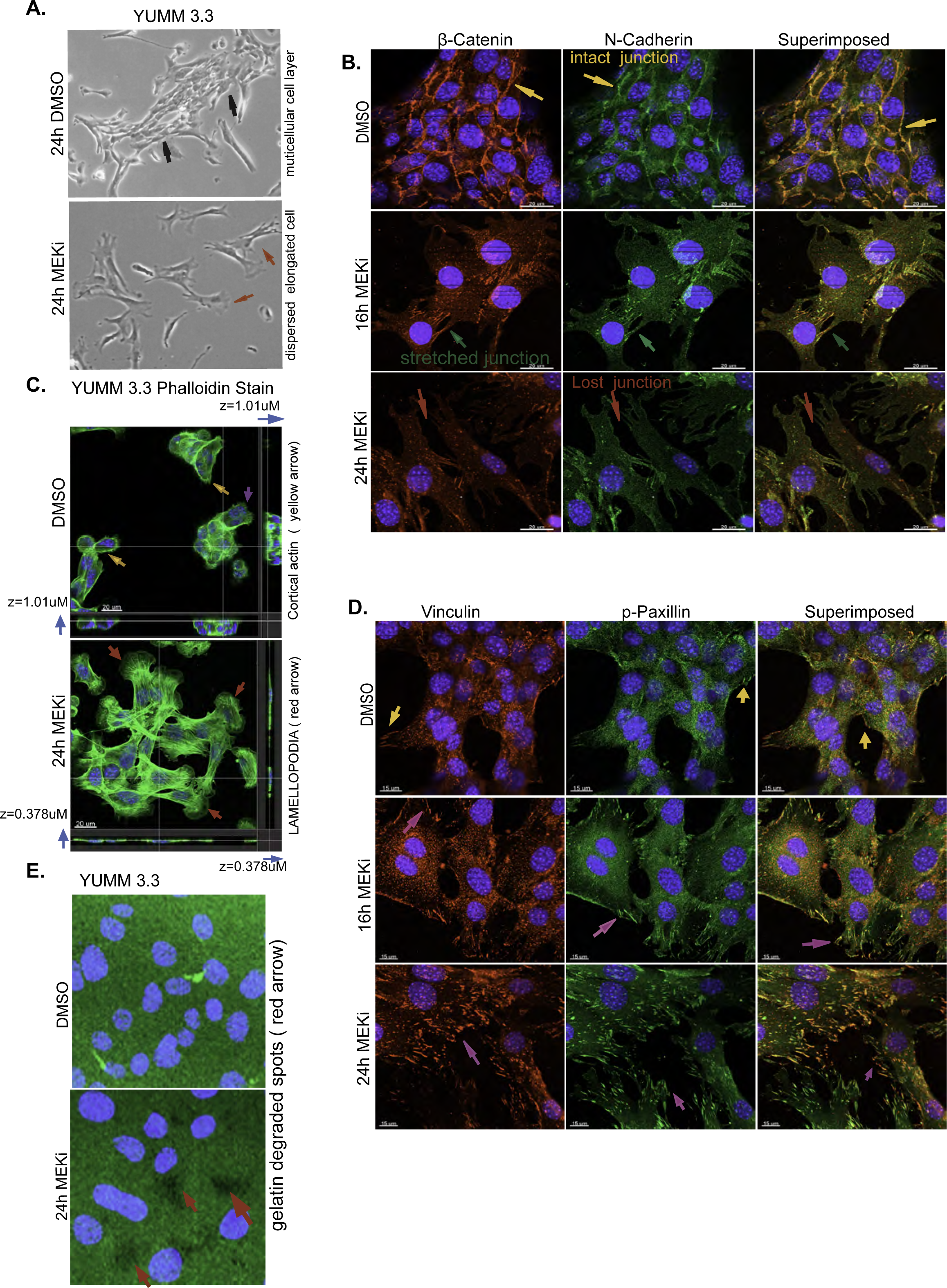
ERK pathway inhibition triggers mesenchymal transition. **A.** Bright-field images of YUMM 3.3 24 hours after treatment with DMSO or 10nM Trametinib. Magnification 10X. **B.** YUMM 3.3 were treated with DMSO or 10nM Trametinib for indicated times, and intercellular junctions were visualized with β-catenin(red) or N-cadherin (green) stains. Superimposed image (orange) shows the co-localization of β-catenin and N-cadherin. Yellow arrow shows intact junctions, green arrow shows stretched junctions, and red arrow shows dissolution of cell junctions. Scale bar, 20uM **C.** In YUMM 3.3, F-actin was visualized with phalloidin (green) after treatment with DMSO or 10nM Trametinib (24h). Images shown are 3D-rendered images of the Confocal Z-stack. Blue arrow shows Z-axis thickness (z), yellow arrow shows cortical actin, and purple and red arrow shows lamellipodia. Scale bar, 20uM. **D.** In YUMM 3.3 focal adhesions were visualized with Vinculin (red) or phospho-Paxillin (Y118) (green) stain after treatment with 10nM Trametinib for indicated times. Superimposed image (orange) shows co-localization of Vinculin and phospho-Paxillin. Purple arrows show focal adhesions dynamics upon ERK inhibition. Scale bar, 20uM. **E.** YUMM 3.3 cells were seeded on plates coated with GFP-labelled gelatin and treated with DMSO or 10nM Trametinib. Twenty-four hours after treatment, gelatin degradation was visualized using a fluorescence microscope. Green channel is gelatin, blue is DAPI, and arrowheads point to spots(black) where gelatin has been degraded. Magnification 40X Representative images of 3 independent experiments are shown. Nuclei were stained with DAPI. Also see SF2, S video 1a, S video 1b, S video 2

Dissolution or weakening of intercellular junctions is a defining feature of the mesenchymal transition. YUMM 3.3 cells were exposed to Trametinib and stained for N-cadherin and β-catenin, major components of intercellular junctions in cells that originate from the neural crest. In DMSO-treated YUMM 3.3 cells, β-catenin (red) and N-cadherin (green) co-localized (orange superimposed) at the cell membrane, thus defining the borders of the adjacent cells and revealing intact intercellular junctions (yellow arrow, **Figure 3B**). Within sixteen hours of Trametinib treatment, cells began to disperse, as shown by the appearance of overstretched intercellular junctions (green arrow). By 24 hours, intercellular junctions were lost, as demonstrated by loss of N-cadherin and Beta-catenin staining at the cell boundary (red arrow). These changes are consistent with the loss of multicellular units of cells and their replacement with elongated single cells with front-back polarity.

Actin treadmilling allows the cell to form lamellipodia which drive the forward movement of mesenchymal cells [22, 23]. The actin cytoskeleton was visualized with phalloidin(green), a peptide that selectively binds to filamentous actin (F-actin). YUMM 3.3 cells treated with DMSO contained a sub-cortical actin mesh (yellow arrow, **Figure 3C),** and only cells with a free edge formed noticeable actin protrusions (purple arrow, **Figure 3C).** After 24 hours of MEK inhibition, gross changes in the actin cytoskeleton had occurred, dominated by the appearance of actin stress fibers traversing the cells and prominent lamellipodia [24, 25] at the leading edge (red arrow, **Figure 3C**). Cell spreading was also apparent, as seen by the reduction in Z-axis thickness(z) (blue arrow), which fell from 1.01uM to 0.378uM with treatment (**Figure 3C, S video 1a, 1b)**. We observed protrusive actin structures (red arrow, **Figure S3A)** and cell spreading in other YUMM and human melanoma models treated with Trametinib. In migrating mesenchymal cells, stress fibers attached to focal adhesions generate the mechanical force necessary for movement [26–28]. In YUMM 3.3 and 3.2, stress fibers began to develop after 8 hours of MEK inhibition and became prominent by 16 hours (red arrow, **Figure S3B)**. We confirmed this in BRAF^V600E^ melanoma models SK-MEL190 and SK-MEL 207(red arrow, **Figure S3C).** A time-lapse video of SK-MEL 207 **(S Video 2)** (follow magenta or blue arrow) with and without (follow red arrow) MEK inhibitor treatment confirms that upon ERK inhibition, cells spread and elongate, forming lamellipodia at the leading edge and pushing the cell forward.

Upon spreading, the leading edge of the cell establishes contact with the extracellular matrix (ECM) by initiating focal adhesion formation. Translocation of the structural protein Vinculin to the focal adhesion stabilizes the focal adhesion complex and strengthens the link to actin filaments, while phosphorylation of the scaffold protein Paxillin allows it to recruit structural proteins and signaling molecules to the focal adhesion complex [29, 30]. We visualized focal adhesion remodeling by staining for Vinculin (green) and Phospho-Paxillin (red) (**Figure 3D**). DMSO-treated YUMM 3.3 cells formed a monolayer devoid of focal adhesions except at the free edges. Focal adhesions were visualized as green, red, or orange (superimposed) dots only at the free border (yellow arrow). By contrast, within 16 hours of MEK inhibition, loosely scattered individual cells acquired green, red, and orange dots, which became prominent by 24 hours (magenta arrow). These data demonstrate that upon MEK treatment, cells spread, and individual cells remodel actin cytoskeleton and focal adhesions, which provide the adhesion and mechanical support for the mesenchymal movement.

Localized proteolytic degradation of extracellular matrix (ECM) is necessary for mesenchymal cell invasion and tissue transmigration [31]. YUMM 3.3 cells were unable to degrade gelatin, as seen by the intact green surface of GFP-coated gelatin in a gelatin degradation assay. After 24 hours of MEK inhibition, we observed degradation of GFP-tagged gelatin as seen by the appearance of black patches devoid of GFP (red arrow, **Figure 3E**), confirming enhanced ECM degrading capacity of MEK inhibitor-treated YUMM 3.3 cells.

Thus, cells with BRAF^V600E^ activation that do not migrate adhere to other cells by means of intercellular junctions and lack actin stress fibers, observable lamellipodia, focal adhesions, and proteolytic activity that would support the mesenchymal migration. Inhibition of ERK signaling leads to a series of events 8-24 hours later **(Figure S3D, S Video 2,** follow magenta or blue arrow) that constitute the mesenchymal transition—loss of intercellular junctions, cell spreading with prominent lamellipodia, formation of actin stress fibers and focal adhesion remodeling. These changes coincide temporally with the induction of migration and invasion. We conclude that, in these cells, ERK activation suppresses migration by suppressing the mesenchymal phenotype. ERK pathway inhibition relieves this suppression and thus induces migration/invasion.

### RAC1 activation after ERK inhibition induces the migration of BRAF^V600E^ cells

RAC1, a member of the Rho family of GTPases, is a key regulator of mesenchymal migration and phenotypic plasticity of cancer cells [32–34]. In the GTP-bound activated state, RAC1 regulates actin cytoskeleton reorganization, lamellipodia formation, focal adhesion remodeling, and proteolytic degradation of ECM [25, 35, 36]. The morphologic changes induced by MEK inhibition in cells with BRAF^V600E^ resembled those observed when RAC1 is activated [34]. Indeed, in YUMM 3.3 cells, RAC1-GTP increased 1.6-fold 8-16 hours after MEK inhibition and 2.6-fold by 32 hours, while RAC1 protein expression was unaffected (**Figure 4A**). This phenomenon was confirmed in five other models, four BRAF^V600E^ melanoma and one mutant NRAS melanoma (**Figure 4A (ii)**). RAC1 was activated upon MEK inhibition in all five cell lines. In three of them, RAC1-GTP rose to significant levels 8-16 hours after drug addition. In two other cell lines, YUMM 3.2 and 1g.1, RAC1 levels rose within one hour of drug exposure. The reason for this could be lower PTEN levels, as YUMM 3.2 has reduced levels of PTEN (**Figure 1A**)., and 1g.1 is PTEN null. Since the reduction of PTEN expression with a short hairpin (shRNA) increases RAC1-GTP (**Figure 6B**), the increased rapidity of induction of RAC1-GTP may be due to decreased PTEN expression. In YUMM 3.3, induction of RAC1-GTP and induction of migration coincided (**Figure 4A (i)),** suggesting that RAC1 activation may play a role in the induction of migration after ERK signaling.

**Figure 4:**
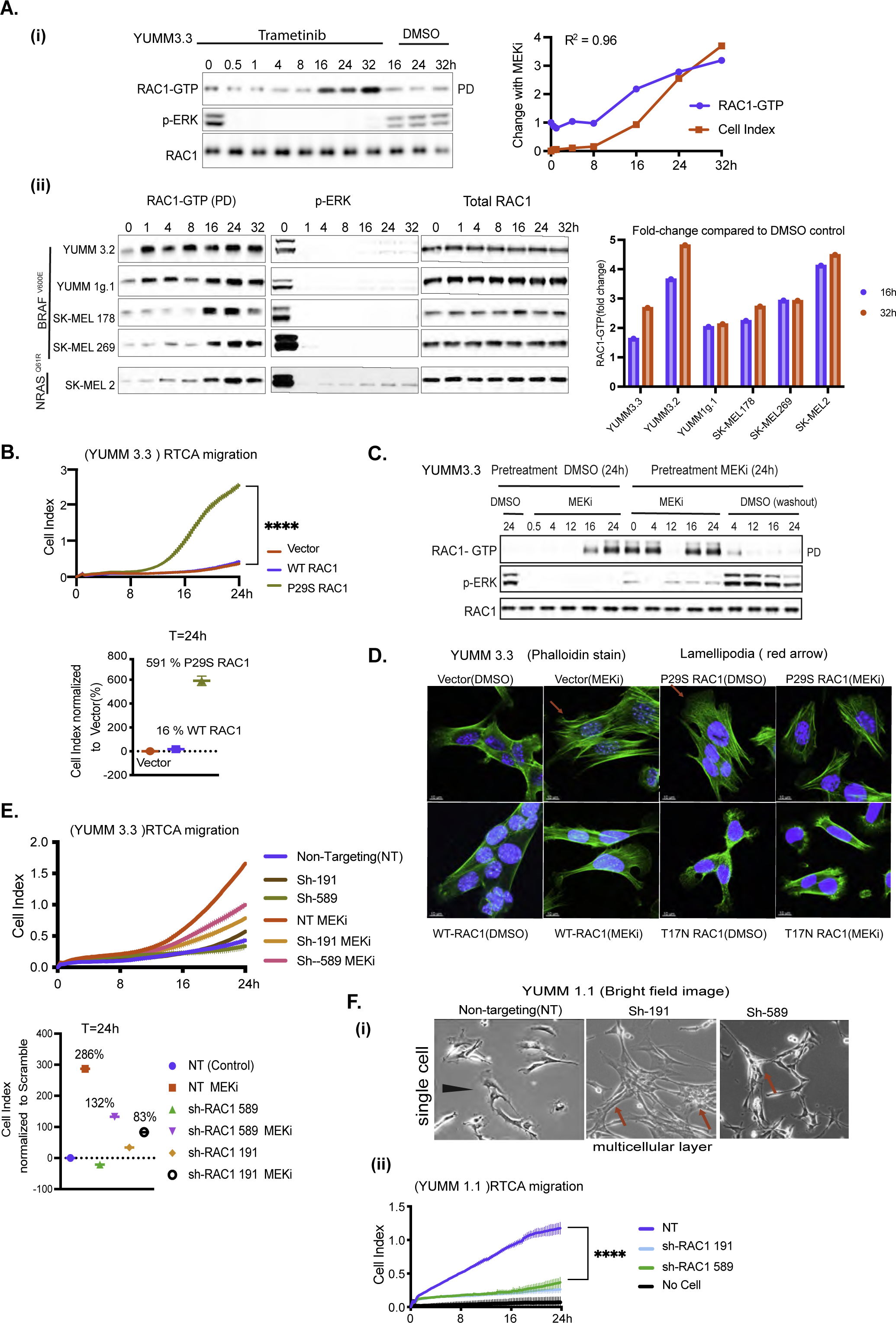
RAC1 activation causes mesenchymal transition and enhances cell motility upon ERK inhibition. **A. i)** YUMM 3.3 cells were treated with 10nM Trametinib (MEKi) for indicated times. RAC1-GTP levels were determined using the active RAC1 pull-down (PD) assay. Total RAC1 and phosphorylated ERK (pERK) was assayed by IB the WCL. Induction of RAC1-GTP and induction of migration upon MEK inhibition were quantified, normalized to DMSO control at T=0h, and plotted as a function of time. (**ii)** Indicated cell lines were treated with Trametinib and RAC1**-**GTP, RAC1 and pERK were detected as above. Bar graph depicts the fold change in RAC1-GTP upon MEK inhibition. **B.** An activated RAC1 mutant rescues migration in BRAF^V600E^ cells. Migration of YUMM 3.3 expressing empty vector, WT RAC1, or constitutively active mutant RAC1^P29S^ (1ug/ml doxycycline 24h). P-values are based on ordinary one-way ANOVA analysis. CelI Index at 24h was normalized to those with an empty vector. **C.** YUMM 3.3 cells were pretreated with DMSO or 10nM Trametinib for 24 hours and each population was split and replated with DMSO or 10nM Trametinib. To wash out the drug, Trametinib pretreated cells were washed with PBS three times before seeding. WCL was collected at indicated times and RAC1-GTP, pERK and Total RAC1 was analyzed as in 4A. **D.** Actin filaments were visualized with phalloidin stain(green) in YUMM 3.3 expressing inducible empty vector, WT RAC1, RAC1^P29S^, or RAC1^T17N^ (1ug/ml doxycycline 24h) after treatment with 10nM Trametinib (24h). **E.** Knockdown of RAC1 expression prevents rescue of migration by MEK inhibition. Migration of YUMM 3.3 cells expressing non-targeting (NT) or two different short hairpins against RAC1: sh-RAC1 19, and sh-RAC1 589, with or without 10nM Trametinib. Cell index at 24h is normalized to NT hairpin with DMSO. P-values are based on ordinary one-way ANOVA analysis **F.** Knock down of RAC1 prevents mesenchymal phenotype and migration **(i)** Brightfield image of YUMM 1.1 cells expressing either NT or short hairpins against RAC1, sh-RAC1 191, and sh-RAC1 589. Magnification 10X **(ii)** Migration of YUMM 1.1 cells expressing either NT or sh-RAC1 191, and sh-RAC1 589. Curve with vertical error bars represents mean ± SEM (n=3). Also, see SF4

To test this hypothesis, we engineered a doxycycline (dox) inducible, constitutively active RAC1 mutant, RAC1^P29S^ into YUMM 3.3 cells [37]. RAC1^P29S^ expression increased RAC1-GTP by 700% **(Figure S4A)** and migration index by approximately 591% compared to empty vector (**Figure 4B**). Overexpression of wildtype RAC1 to levels similar to RAC1^P29S^ expression increased RAC1-GTP by 37% and migration index by 16% at 24 hours. These data support the idea that ERK activation suppresses RAC1 activation, which hinders cell migration, and that induction of RAC1 activation after ERK pathway inhibition is responsible for the induction of migration.

To determine whether induction of RAC1-GTP with MEK inhibitors was reversible, YUMM 3.3 was treated with MEK inhibitor Trametinib or DMSO for 24 hours. At 24 hours, pretreated cells were replated with fresh media containing DMSO or Trametinib. (**Figure 4C).** In DMSO pretreated cells, RAC1-GTP was low, and p-ERK was high. MEK inhibition caused rapid inhibition of p-ERK and, 16 hours later, induction of RAC1-GTP. In MEK inhibitor pretreated cells, p-ERK was low, and RAC1-GTP was elevated. and when MEK inhibitor was re-added, RAC1-GTP remained elevated. But when MEK inhibitor was washed out, p-ERK rapidly rose, and RAC1-GTP fell within 4 hours. Thus, activation of RAC1 by ERK pathway inhibition is reversible.

We asked whether RAC1^P29S^ causes the morphologic changes elicited by ERK inhibition. RAC1^P29S^ expression induced prominent lamellipodia (red arrow, **Figure 4D**), similar to those induced in BRAF^V600E^ cells treated with MEK inhibitor. Cells expressing dominant-negative mutant RAC1^T17N^ didn’t spread or form stress fibers and lacked lamellipodia. Moreover, MEK inhibition failed to reinduce cell spreading or lamellipodia formation in these cells, which is consistent with work showing the importance of activated RAC1 for these processes [38, 39]. Cells with RAC1^P29S^ also formed focal adhesions at cell periphery similar to those formed in YUMM 3.3 cells after MEK inhibition (purple arrow, **Figure S4B)**. Cells with wildtype RAC1 had focal adhesions only at the outer edges of cell patches (yellow arrow), whereas cells with dominant negative mutant RAC1^T17N^ had almost none at the cell borders. These data suggest that RAC1 activation contributes to focal adhesion dynamics upon ERK inhibition. Though RAC1^T17N^ expression was minimal compared to wildtype or RAC1^P29S^ **(Figure S4A),** it inhibited RAC1-GTP by 63% compared to vector, and this was enough to oppose the morphological changes elicited by RAC1 activation.

To confirm the role of RAC1 in the mesenchymal transition and enhanced migration induced upon ERK inhibition, we knocked down RAC1 in YUMM 3.3**(Figure S4C)** and YUMM 1.1 **(Figure S4E)** with two different short hairpins (sh). In 3.3, sh-191 and sh-589 downregulated RAC1 expression to 15% and 59% of baseline with concurrent downregulation of RAC1-GTP. Knocking down RAC1 inhibited the loss of intercellular junctions upon MEK inhibition, as shown by intact beta-catenin stain (blue arrow, **Figure S4D)**. This was accompanied by a blunting of induction of cell migration by the MEK inhibitor (**Figure 4E**). Upon MEK inhibition, migration index was induced by 286% with non-targeting hairpin, 83% with sh-191, and 132% with sh-589.

In YUMM 1.1(BRAF^V600E^ PTEN^null^), a cell line with a prominent population of single cells (black arrow, **Figure 4F, (i))**, RAC1 knockdown caused formation of multicellular units with cell-cell junctions (red arrow), losing its mesenchymal morphology and concomitant inhibition of cell migration (**Figure 4F (ii)).** Knockdown of RAC1 in YUMM 1.1 or blocking RAC1 activation with dominant negative mutant RAC1^T17N^ in SK-MEL 190 cells suppresses ERK inhibition induced migration **(Figures S4F, S4G)**. RAC1^T17N^ also suppressed the phosphorylation of RAC1 substrate PAK1, suggesting effective inhibition of RAC1 activation **(Figure S4G).** Taken together, these data confirm that changes in cell morphology and the increase in migration observed upon ERK pathway inhibition require induction of RAC1-GTP.

### Relief of ERK-dependent feedback inhibition of receptor signaling activates RAC1 and increases the migration of BRAF^V600E^-driven tumors

Our findings suggest that inhibition of RAC1 activation by ERK signaling significantly retards the migration and invasion of BRAF^V600E^ melanomas. BRAF^V600E^-driven ERK signaling is known to cause feedback inhibition of receptor tyrosine kinases (RTKs) signaling [40], an upstream activator of RAC1 [41–44].

Doxycycline-induced expression of BRAF^V600E^ in MEFs inhibited PDGF receptor (PDGFR) and EGF receptor (EGFR) phosphorylation, and this inhibition was rescued by MEK inhibition, suggesting their suppression by ERK (**Figure 5A).** We examined whether inhibition of ERK signaling induces receptor phosphorylation in YUMM 3.3 and human melanoma models. PDGFR phosphorylation was induced in YUMM 3.3 cells 8-12 hours after MEK inhibitor treatment (**Figure 5B**).

**Figure 5:**
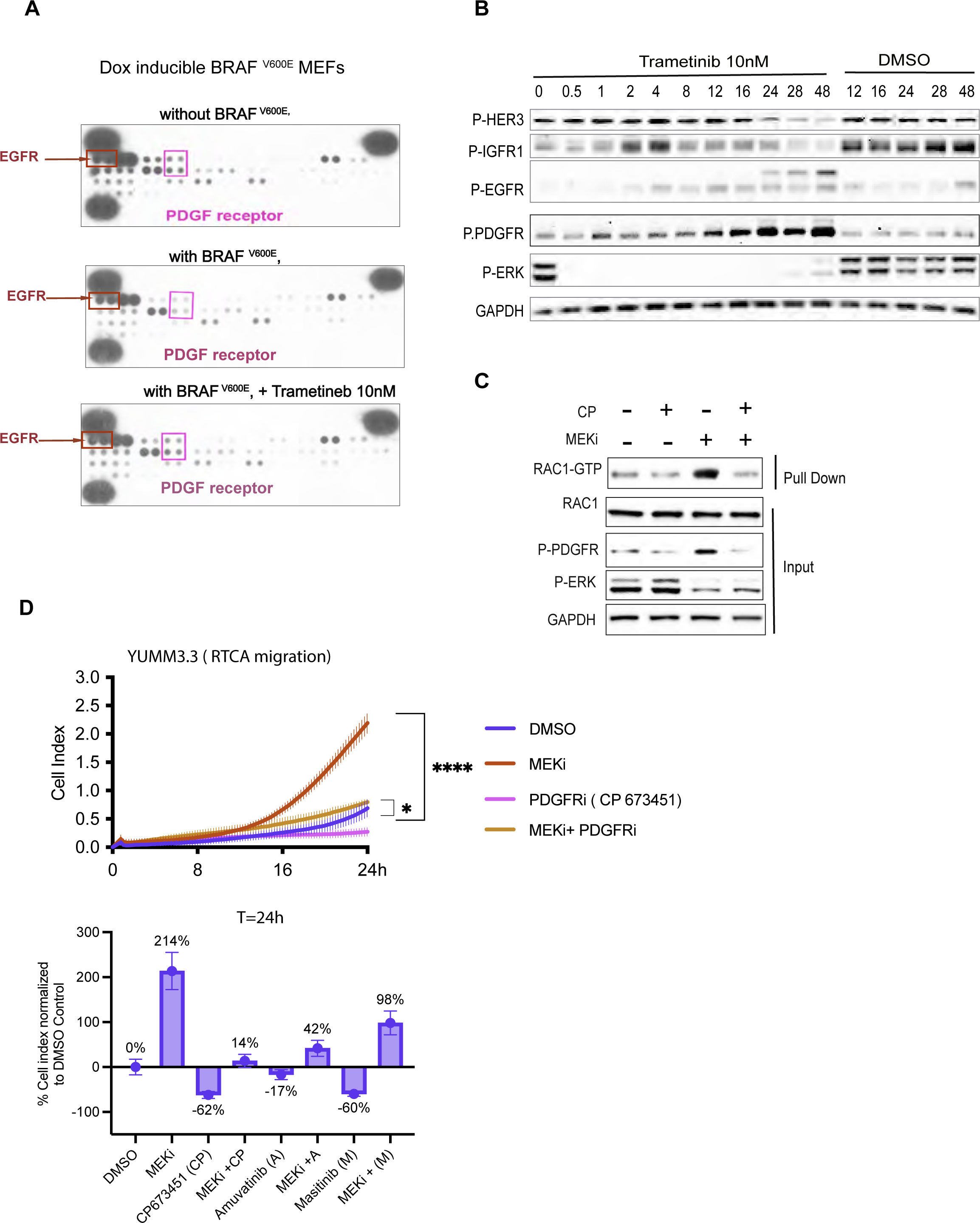
Relief of ERK-dependent feedback potentiates RTK signaling, which in turn activates RAC1 and cell migration. **A.** MEFs expressing BRAF^V600E^ (1ug/ml doxycycline 24h) were treated with Trametinib for 24h, and WCL was collected. WCLs were analyzed by an RTK array to profile phosphorylated RTKs. MEFs with empty vector was used as control. **B.** YUMM 3.3 were treated with 10nm Trametinib for indicated times, and WCL was IB for indicated proteins. **C.** YUMM 3.3 were pretreated with 1uM PDGF receptor inhibitor CP673451(CP) for 1 hour followed by DSMO or 10nM Trametinib (24h). WCL was collected and subjected to RAC1 PD assay or IB for indicated proteins. **D.** Migration of YUMM 3.3 cells treated with DMSO, 10 nM Trametinib, and 1uM CP alone or in combination with Trametinib. Cell Index at 24 hours is normalized to DMSO control and plotted as a bar graph for all three PDGFR inhibitors used. Vertical error bars on curve represents mean ± SEM (n=3). Also, see SF5

Phosphorylation of HER3, EGF and IGF receptors was also induced, with different kinetics. Upon ERK inhibition EGFR, HER2, HER3, and PDGFR phosphorylation were also induced in SK-MEL 190 and A375 **(Figure S5A**). In YUMM 3.3, induction of RAC1-GTP, PDGFR phosphorylation, and increased migration upon MEK inhibitor treatment all begin 8-16 hours after drug addition. The upregulation of total receptor expression accompanied the increase in receptor phosphorylation in all three cell lines **(Figure S5A).**

In YUMM 3.3 cells, baseline RAC1-GTP and its induction by MEK inhibition were both reduced by pretreatment with the selective PDGFR kinase inhibitor CP-673451 (CP) (**Figure 5C).** PDGFR inhibition by three different inhibitors, CP673451, Amuvatinib (A), and Masitinib(M) (**Figures 5D, S5B),** reduced induction of migration by MEK from 214% to 14%, 42% and 98% of DMSO control respectively without affecting cell viability.

Taken together, these results suggest that ERK-dependent feedback inhibition of PDGFR and perhaps other RTKs impair the migration of melanocytes expressing BRAF^V600E^ by inhibiting RAC1 activation. Moreover, relief of ERK-dependent feedback inhibition of receptors by ERK pathway inhibition restores migration. These conclusions are supported by our demonstration in our systems that RAC1 activation is regulated by RTK signaling, and RAC1-GTP is required for migration.

### BRAF^V600E^ melanomas harbor mutations that rescue cell migration and invasion

Our data support the idea that BRAF^V600E^ suppresses cell migration and invasion by causing ERK-dependent feedback inhibition of RAC1 activation. We, therefore, hypothesize that secondary hits that restore migration are required for melanomagenesis. Our data imply that this can be accomplished by lesions that cause RAC1 activation to be insensitive to ERK-dependent feedback inhibition. In support of this hypothesis, *RAC1* mutations (7%), amplification or mutations of PI3K-dependent RAC exchange factors 1 and 2 (*PREX1, PREX2)* (*31*%), and lesions that inactivate *PTEN* (15%) were identified in a set of 287 human melanomas (TCGA Firehose legacy, **Figure 6A).** Most of the RAC1 mutants identified in melanoma are gain-of-function P29S mutants that we show here rescue migration and the mesenchymal phenotype in BRAF^V600^ expressing melanocytes (**Figure 4B**). PREX1 and PREX2 are PI3K-dependent, RAC1 nucleotide exchange factors that switch RAC1 to active GTP-bound conformation. PREX proteins are activated by PI3K and phosphorylated and inhibited by RAC1-activated PAK kinases [45, 46]. PTEN is thought to inhibit RAC1 activation by inhibiting PI3K signaling [47]. In melanomas, *PTEN* lesions have been associated with loss of protein expression.

**Figure 6:**
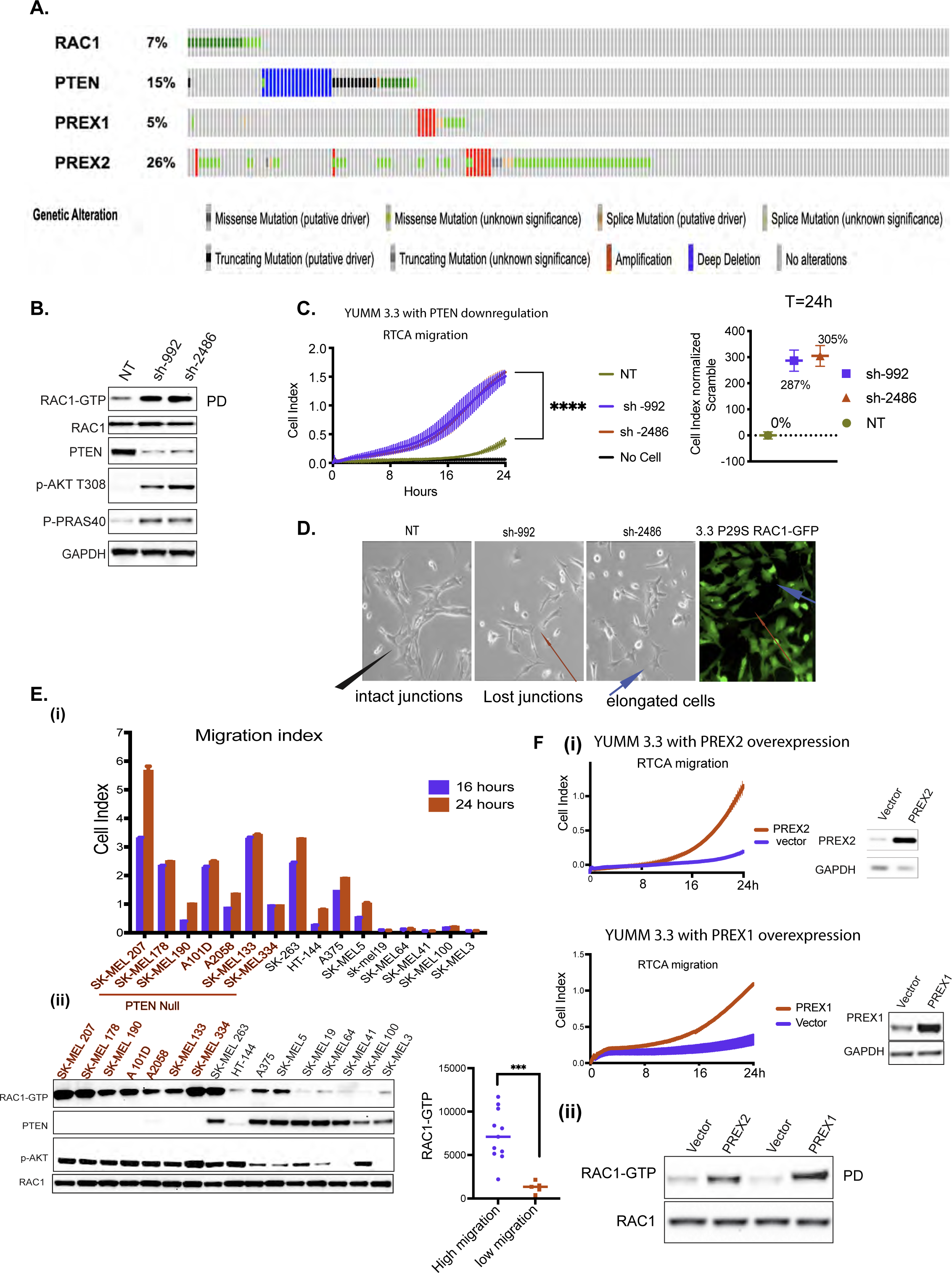
BRAF^V600E^ melanoma is enriched with lesions that rescue cell motility. **A.** Oncoprint of melanoma patients with lesions in *PTEN, RAC1, PREX1,* or *PREX2* genes obtained from Firehose legacy genomics dataset from MSKCC cBioPortal for Cancer Genomics. **B.** YUMM 3.3 expressing either NT or two different short hairpins against PTEN, sh-992, and sh-2486 was assayed for RAC1-GTP and RAC1 expression as in Fig 4A. WCLs were also IB with indicated antibodies. **C.** Migration of YUMM 3.3 expressing NT or short hairpins against PTEN, sh-992, and sh-2486. Curve with vertical error bars represents mean ± SEM (n=3). Cell index at 24h is normalized to control (NT) with DMSO. **D.** Brightfield image of YUMM 3.3 with PTEN KD or RAC1^P29S^ expression. Black arrow in control (NT) shows multicellular unit, and red arrow shows lost junctions upon PTEN KD. GFP tagged RAC1^P29S^ was imaged with a fluorescence microscope to show RAC1^P29S^ expression. (Blue arrow shows actin protrusions). Magnification 10X **E.** Cell index of indicated cell lines at 24h and 16h. RAC1-GTP was assessed as describe Fig 4A. WCL was assayed by IB with the indicated antibodies. RAC1-GTP was quantified, and it’s distribution was plotted against cells migration indexes (cell index at 24h). **F. i)** Migration of YUMM 3.3 cells overexpressing PREX1 or PREX2 (transient transfection 36h). PREX1 and PREX2 expression was assayed by IB. **ii)** WCLs were assessed for RAC1-GTP and RAC1 as in 4A. Also, see SF6

Thus, RTK-PI3K(PTEN)-PREX1,2-RAC1-PAK-kinases constitute a potential pathway that regulates RAC1 function and migration. These lesions occur in 53% of melanomas (**Figure 6A).** We hypothesize that, like RAC1^P29S,^ these lesions function at least in part by activating RAC1 and stimulating the migration of BRAF^V600E^ melanocytes. In YUMM 3.3 downregulation of PTEN expression with two different hairpin*s*, sh-992 and sh-2486, caused 3.1 and 3.8-fold induction of RAC1 GTP (**Figures 6B, S6A)** and a concomitant increase in migration by 287% and 305% respectively, compared to controls (**Figure 6C**). These data are consistent with those in (**Figures 1A, S1B, S1C)** in which cells with co-existent PTEN loss and BRAF^V600E^ migrate and invade the ECM faster than those with BRAF^V600E^ alone. PTEN downregulation was also associated with a change in cell morphology similar to that caused by RAC1^P29S^ (red arrow, **Figure 6D**). Upon PTEN knockdown, morphology of YUMM 3.3 changed from a multicellular layer to a mixed population of clustered or elongated cells with actin protrusions or lamellipodia (blue arrow). YUMM 1.1 (PTEN^null^) cells are elongated single cells (blue arrow) as opposed to multicellular patches of YUMM 3.3 (PTEN^WT^) (red arrow, **Figure S6B)**. Moreover, RAC1 activation is required for the migration of BRAF^V600E^ PTEN^null^ cells. The expression of the dominant negative RAC1^T17N^ mutant inhibited migration of BRAF^V600E^ PTEN^null^ YUMM1.1 and SK-MEL 190 cells **(Figure S6C).**

We assessed the migration of 16 human melanoma lines with BRAF^V600E^. Five of these had significantly lower migration indices than the others (**Figure 6E i**) and lower levels of RAC1-GTP (**Figure 6E ii).** We determined these cell lines all had wildtype PTEN, RAC1, PREX1, and PREX2 (cbioportal, data not shown) and had low levels of p-AKT compared to migratory lines. The other 11 lines migrated significantly faster. Of these, 8 were PTEN^null,^ SK-MEL 263 had wildtype PTEN and RAC1 amplification (cbioportal, data not shown), and elevated RAC1-GTP. There were three outliers with moderate migratory ability compared to PTEN^null^ cells. HT-144 was haploinsufficient for PTEN, with high p-AKT but low RAC1-GTP. A375 and SK-MEL5 had wild-type PTEN, low p-AKT levels, moderate RAC1-GTP levels, and moderate migratory ability compared to PTEN^null^ cells. In this cohort of cells, cell migration positively correlates with RAC1-GTP levels **(graph Figure 6E ii).** 5% of melanoma have lesions in *PREX1,* and 26% have lesions in *PREX2* genes (**Figure 6A**) [48, 49]. *PREX1* or *PREX2* are amplified or have missense mutations, most of which are not characterized. Some characterized truncating PREX2 mutants, such as E824* and K278*, have increased RAC1 GEF activity [50]. As expected from their function, we found PREX1 or PREX2 overexpression induced RAC1-GTP and rescued cell migration in YUMM 3.3 (**Figure 6F).** These data support the idea that RTK-activated pathway comprised of an RTK, PI3K, PREX1 or PREX2, and RAC1 regulate RAC1 activation, and those genetic events that activate components of the pathway are sufficient to bypass the feedback inhibition of RAC1 by BRAF^V600E^ and induce migration of BRAF^V600E^ expressing melanocytes.

## Discussion

The kinase activity of BRAF^V600^ mutants is hyperactivated and drives elevated ERK signaling output accompanied by exaggerated ERK-dependent feedback inhibition of upstream signaling, including that of RTKs and WT RAS-activated pathways. However, BRAF^V600^ is not sufficient to cause common human tumors. In mouse models, it has been shown that BRAF^V600E^ causes benign melanocytic hyperplasia followed by permanent growth arrest. However, disseminated melanoma rapidly develops in GEMMs engineered to express BRAF^V600E^ in melanocytes with co-existent PTEN loss [9–11].

We set out to understand this phenomenon. BRAF^V600E^ PTEN^null^ melanocytes did not grow more rapidly than those with BRAF^V600E^ alone. Instead, our findings suggest that melanocytes expressing BRAF^V600E^ do not form malignant disease because, although high levels of ERK signaling deregulate proliferation, they also feedback inhibit the RAC1 GTPase, which is required for mesenchymal phenotype and cell migration. PTEN loss rescues RAC1 activation, leading to mesenchymal transition and enhanced migration, enabling malignant transformation.

This model is supported by the inhibition of migration by BRAF^V600E^ in MEFs, acceleration of migration, and invasion of BRAF^V600E^ cells after inhibition of ERK signaling in culture and *in vivo*. Enhanced migration upon ERK inhibition is associated with the induction of a mesenchymal phenotype consistent with the activation of RAC1 and mesenchymal migration. Consistent with these data, MEK inhibition has been shown to increase the invasive behavior of BRAF^V600E^ melanoma cells into 3D fibrillar collagen [53]. The mechanism by which ERK inhibition activates RAC1 is the release of negative feedback of PDGFR. Inhibition of PDGFR inhibits the induction of RAC1-GTP and migration caused by MEK inhibition. Inhibition of RAC1 by expression of a dominant negative RAC1 mutant or by knockdown of RAC1 expression prevents both the mesenchymal transition and migration that occur with MEK inhibition. Thus, feedback inhibition of PDGFR is responsible for suppressing RAC1-GTP and cell migration by BRAF^V600E^ in our model system. The feedback network is complex, and feedback inhibition of other upstream RTKs likely plays a role in other tumors. RAS also activates ERK, but tumors with mutant RAS do migrate. This is likely due to other RAS effectors—PI3K and TIAMs—that activate invasion and RAC1. In fact, although tumor cells with mutant RAS migrate, we show that MEK inhibition increases migration. Thus, RAS activates a balance of inhibitors and stimulators of migration. Control of the effectors may play a role in regulating the mesenchymal switch.

These results are consistent with work of the Downward laboratory. [15] which showed that neither RAF mutants, PI3K mutants, nor RAS mutants that activated RAF but not PI3K could fully transform murine fibroblasts. However, these RAF-activating RAS mutants synergized with RAS mutants that activate PI3K or vectors that drive activated RAC1 or PI3K. Moreover, RAS activation of PI3K or RAC1, but not ERK, was required for actin remodeling and stimulating membrane ruffles. Activated BRAF mutants that activate ERK and not PI3K had not yet been identified in human tumors, but their work predicts that if such mutants existed, they would stimulate proliferation but not induce membrane ruffling or actin reorganization. We now show that BRAF^V600E^ causes ERK-dependent feedback inhibition of upstream receptor signaling and, by this mechanism, inhibits activation of WT RAC1.

We conclude that, whereas hyperactivation of ERK signaling by oncogenic BRAF mutants is sufficient to deregulate cell proliferation, ERK-dependent feedback inhibition of RAC1 suppresses migration. Secondary mutations that enhance migration in BRAF mutant melanocytes despite this feedback are thus necessary for tumor formation. Our data support the concept that one potential consequence of oncoprotein driven feedback is the inhibition of processes required for transformation. In that case, secondary events that rescue that process would be required for tumorigenesis. We have identified four lesions that co-occur with BRAF^V600E^ melanoma—PTEN loss, PREX1 and 2 amplification or mutation, and a RAC1 activating mutation-each of which rescues RAC1 activation and migration. At least one of these lesions occurs in approximately 48% of melanomas. These lesions may have multiple other advantages for tumor progression, but our data confirms that one function they serve is to rescue cell migration and invasion.

Consistent with our hypothesis, constitutively activated RAC1^P29S^ mutant in combination with mutant BRAF has an oncogenic function in melanoma [51]. We believe this oncogenic function is due to its RTK and RAS-independent induction of the mesenchymal phenotype that is resistant to ERK-dependent feedback.

The model explains the selection of these secondary mutants, the poor transforming ability of BRAF^V600E^ by itself, the high frequency of BRAF^V600E^ in benign nevi that only rarely undergo transformation, and the slow-growing, non-invasive phenotype of pediatric brain tumors in which RAF mutants or fusions are the lone mutations [52]. It also may contribute to the oft-cited inverse relationship between proliferation and migration. ERK activation may inhibit migration while driving proliferation.

Contrary to our data, others have shown that BRAF^V600E^ cells do migrate well. It is possible that these cells contain other mutations that overcome feedback inhibition of RAC1 or activate migration by other mechanisms.

Therefore, we believe we have discovered a new negative function of oncogenes: feedback inhibition of processes required for transformation that must be rescued by other mutations for tumors to develop and evolve. We have identified one, but given the ubiquity of feedback mechanisms, there are likely to be others.

## Acknowledgements

We are grateful to Mahesh Saqcena (James Fagin Laboratory MSKCC) for sharing BRAFV600E Thyroid cells and to Sebastian Carrasco (MSKCC) for reviewing the histological slides and for his helpful discussion.

## Funding

This research was supported by grants (to N.R.) from the National Institutes of Health (NIH) P01-CA129243; R35 CA210085; the Geoffrey Beene Cancer Research Center; the Emerson Collective Research Grant; Melanoma Research Alliance; The NIH MSKCC Cancer Center Core Grant P30 CA008748 and Experimental Therapeutics Center. S.F.R is funded by a grant from Astra Zeneca (FE 10050024 Yale SRA), the Melanoma Research Alliance (1154713), the Canadian Institute of Health Research (CIHR 490134), and the Fonds de Recherche du Québec en Santé (FRQS 329696).

## Authors Contributions

S.G., and N.R. conceived the hypotheses, and wrote the manuscript. N.R., M.B., S.G., J.B., E.S., K.T., M.O., and R.Y., contributed to experimental design and data analysis. S.G, J.B, N.O., M.S., S.F.R., H.L, N.F., E.R., Y.R., A.B., Q.C., P.P., N.T., performed experiments

## Declaration of interests

(Authors declare no potential conflicts of interest) N.R. is on the scientific advisory board (SAB) and owns equity in Beigene, Zai Labs, MapKure, Ribon and Effector. N.R. is also on the SAB of Astra Zeneca and Chugai and a past SAB member of Novartis, Millennium-Takeda, Kura, and Araxes. N.R. is a consultant to RevMed, Tarveda, Array-Pfizer, Boeringher-Ingelheim and Eli Lilly. He receives research funding from Revmed, AstraZeneca, Array Pfizer and Boerhinger-Ingelheim and owns equity in Kura Oncology and Fortress. M.B has received research funding from Astra Zeneca. R.Y. has served as an advisor for Amgen, Array BioPharma/Pfizer, Mirati Therapeutics, and Zai Lab and has received research support to her institution from Array BioPharma/Pfizer, Boehringer Ingelheim, Daiichi Sankyo, and Mirati Therapeutics. S.F.R is funded by a grant from Astra Zeneca (FE 10050024 Yale SRA), the Melanoma Research Alliance (1154713), the Canadian Institute of Health Research (CIHR 490134) and the Fonds de Recherche du Québec en Santé (FRQS 329696).

## METHODS

### Cell Lines and Compounds

YUMM 4.1 (RRID: CVCL _ JK38), YUMM 1.3 (RRID: CVCL _ JK12), YUMM 1.1 (RRID: CVCL_JK10), YUMM 1.G1 (RRID: CVCL _ JK25), YUMM 3.2 (RRID: CVCL _ JK35), YUMM 3.3 (RRID: CVCL _ JK36), YUMM 3.3 expressing GFP (YUMM 3.3 derived cell line), and Mouse embryonic fibroblasts (MEFs) (ATCC, CRL-2991) were maintained in DMEM/F12 (Corning). All YUMM lines were provided by Dr Marcus Bosenberg’s laboratory at Yale University. SK-MEL 207, A375, SK-MEL 190, SK-MEL 269, SK-MEL 2, WM3918, SK-MEL 178, A101D, A2058, SK-MEL133, SK-MEL 334, SK-MEL 263, HT-144, SK-MEL5, SK-MEL19, SK-MEL3, SK-MEL 100, SK-MEL41, SK_MEL64 were maintained in RPMI 1640 (Corning). B47275 was maintained in Coon’s/F12 (Corning). B47275 cell line was kindly provided by Dr James Fagin Laboratory MSKCC. YUMM 3.3 derived cell lines YUMM 3.3 sh-RAC1 191, YUMM 3.3 sh-RAC1 589, YUMM 3.3 sh-Scramble, YUMM 3.3 sh-PTEN 992, YUMM 3.3 sh-PTEN 2486 and YUMM 1.1 derived cell lines YUMM 1.1 sh-RAC1 191, YUMM 1.1 sh-RAC1 589, and YUMM 1.1 sh-Scramble were maintained in DMEM/F12 supplemented with 0.5ug/ml puromycin (ThermoFisher Scientific, A1113802). YUMM 3.3 derived cell lines YUMM 3.3 empty vector, YUMM 3.3 WT RAC1, YUMM 3.3 T17N RAC1 and YUMM 3.3 P29S RAC1 and YUMM 1.1 derived cell lines YUMM 1.1 empty vector, YUMM 1.1 WT RAC1, and YUMM 1.1 T17N RAC1 were maintained in DMEM/F12 supplemented with 10% tetracycline free FBS(Corning), 2 mM L-glutamine (Corning), 20 units/ml penicillin, and 20 mg/ml streptomycin (Corning), 0.5ug/ml puromycin and 200ug/ml Hygromycin (ThermoFisher Scientific, 10687010). All cell lines were maintained in a humidified atmosphere with 5% CO2 at 37 **o**C. SK-MEL 190 derived cell lines SK-MEL190 empty vector, SK-MEL190 wildtype RAC1, and SK-MEL190 T17N RAC1 were maintained in RPMI 1640 supplemented with 10% tetracycline free FBS, 2 mM L-glutamine, 20 units/ml penicillin, and 20 mg/ml streptomycin, 0.5ug/ml puromycin and 200ug/ml Hygromycin. All media were supplemented with 10% FBS, 2 mM L-glutamine, 20 units/ml penicillin, and 20 mg/ml streptomycin unless mentioned otherwise.

PLX4720 and Vemurafenib was provided by Plexxikon, Trametinib was provided by GlaxoSmithKline, SCH772984 (#S7101), Amuvatinib (#S1244), Masitinib (#S1064) and CP 673451 (#S1536) was purchased from Selleckchem.

### RTCA cell migration and invasion assay

RTCA instrument (ACEA Biosciences) used to monitor cell migration or cell invasion in real-time was maintained in a humidified atmosphere with 5% CO2 at 37°C. For migration or invasion assay, cells were split 24 hours before the assay at 80% confluence. On the day of the assay, 160ul of regular media (media in which the cells are maintained regularly) was added to the lower chambers of CIM-plate 16(Fisher Scientific, #NC0552303), and the upper chamber was attached to the lower chambers. For migration assay, 30ul of regular media was added to the upper chamber, and the assembled CIM-plate was placed in the RTCA analyzer for 1 hour. After one hour, baseline measurements were taken. For invasion assay, upper chamber was coated with 30ul Matrigel (Corning, # CLS356234) (dilution 1:40 diluted in cell media). The assembled CIM-plates were placed in RTCA for 4 hours. After 4 hours, baseline measurements were taken. After taking the baseline measurement, 80,000 cells in 90ul of the regular media/well were added to the upper chamber. In addition, in the control samples, 40ul of DMSO-containing media was added, while samples receiving treatment received 40ul of media supplemented with the drug. Cells were allowed to settle for 15 minutes, the CIM plate was placed on the RTCA chamber, and the Cell Index was recorded every 15 minutes. When the assay was complete Cell Index was analyzed and plotted as a function of time using Graph-pad prism. All assays were performed with a minimum of three replicates and were each repeated three times.

### Transwell cell migration and invasion assay

For transwell cell migration or invasion assay, cells were split at 80% confluence 24 hours before the assay. On the day of the assay,750ul of regular media (media in which the cells are maintained regularly) was added to the bottom chambers of the transwell plates, and 8μm pore-sized transwell filters (Corning, #3422) were inserted into the bottom chambers. For migration assay, 30ul of regular media was added to the transwell filters and the transwell plate was placed in an incubator for an hour. For invasion assay, transwell filters were coated with 30ul Matrigel (Corning, #CLS356234) (dilution 1:40 diluted in cell media) and the assembled transwell plate was placed in an incubator for 4 hours. After placing the assembled transwell plate for an hour (migration assay) or 4 hours (invasion assay) in the incubator, 80,000 cells in 90ul of the regular media/ well were seeded into the filters. In addition, in the control samples, 40ul of DMSO-containing media was added, while samples receiving treatment received 40ul of media with drugs and the assembled transwell plate was placed back in the incubator. At the defined time (endpoint), the filters were fixed with 4% PFA (Fisher Scientific, #50-980-495), and the underbelly of the filter was stained with 0.2% crystal violet (Sigma Aldrich, #C0775-25G). Each testing group contained at least three independent wells. Digital image of the filters was taken using Zeiss AxioImager.Z2, and Fiji software was used to determine the crystal violet positive area from those images.

### Scratch assay

For the wound healing assay, cells were seeded in 10 cm plates with either DMSO or drug at 90% confluence and incubated for 24 hours. Scratch was made with 10ul pipette tip at 24 hours. Brightfield images of the wound area were taken, and the area used was marked. Cells were allowed to migrate, and the Bright field image of the same wound area was taken after 20 hours of wounding using EVOS Brightfield microscope. Fiji software (RRID: SCR_002285) was used to calculate wound area from those images. Images of three independent wound areas were taken, and each assay was repeated three times.

### Cell Growth Assay

Cells were seeded into 96-well plates at 1200 cells per well. After 24 hours, cells were treated with either DMSO or indicated concentration of drugs. Cell growth was measured using the CellTiter-Glo® 2.0 Cell Viability Assay (Promega,# G9242) at each indicated time point. Assays were performed with three independent replicates, and each experiment repeated at least three times. In each experiment, GraphPad Prism was used to generate cell growth plots.

### Immunoblotting

Cells were lysed in RIPA buffer (Thermo Fisher Scientific, # 89901) supplemented with protease and phosphatase inhibitors (Pierce, #87786, #78420). Lysates were centrifuged at maximum speed for 15 min, and protein concentration was determined using the BCA kit (Fisher Scientific, #23225). Samples were prepared using LDS sample buffer (Invitrogen, #NP0008) + Reducing agent (Invitrogen, #NP0009). 10-30ug of protein lysate was loaded onto 4–12% Bis-Tris gels (Thermo Fisher, # NP0321BOX) followed by transfer to nitrocellulose membrane (Bio-Rad, # 1620233). Membranes were incubated overnight with indicated primary antibodies, followed by incubation with secondary antibodies, and developed using iBrightCl1000. Detection was performed using Immobilion Western (Millipore, # WBKLS05000), or West Femto Substrate (Thermo Scientific, # 34096). Primary antibody was used at a concentration of 1:1000. Anti p-ERK1/2 (#4370), anti-Total-ERK1/2 (#4695), anti-p-MEK1/2 (#9154), anti-p-90 RSK (#9341), anti-p-PAK (# 2601), anti-p-MEK S298 (# 2198), anti-p-PDGFR (# 4549), anti-Total-PDGFR (# 3169), anti-p-HER2 (# 2247), anti-p-HER3(# 4791), anti-Total-HER3 (# 4754), anti-p-EGFR (# 3777), anti-Total-EGFR (# 2232), anti-p-IGFR1(# 3918), anti-p-AKT (# 4060), anti-p-PRAS40 (# 13168), anti-PREX1 (# 13168), and PTEN (# 9188) were purchased from Cell Signaling. Anti-GAPDH (sc-47724), anti-BRAF (sc-5284), and anti-CyclinD1(sc-8396) were obtained from Santa Cruz Biotechnology. Anti-PREX2 (# PA5-117672) and anti-RAC1 (#16118) were obtained from Thermo Scientific. anti-Flag (# F1804), and Goat anti-Rabbit secondary (# A4914) was purchased from Sigma. Sheep-anti-mouse secondary (# NXA931) was purchased from GE Healthcare.

### In vivo invasion assay

#### Cell preparation

YUMM 3.3 cells expressing GFP, and luciferase were split 24 hours before the injection and plated at 70% confluence in the presence of either DMSO or 1uM PLX4720. On the day of injection, cells were trypsinized, and trypsin was deactivated with media used to maintain YUMM 3.3. Cells were washed twice with sterile 1× PBS containing PLX4720 or DMSO and counted.

#### Intradermal injection

100,000 cells in 50:50 Matrigel (Corning) were injected intradermally in the flank of six-to eight-week-old female C57BL/6J mice (RRID:IMSR_JAX:000664). Injection sites were marked with tissue marking ink. 24 hours post-injection, treatment was initiated with either a control vehicle or PLX4720 (Plexxikon, Berkeley, CA) at a dose of 7.5 mg/kg twice daily via oral gavage. Mice were randomized (n=3 mice per group) to receive drug treatment or vehicle as a control. Tumor volumes and weights were measured twice a week.

Mice were monitored for the appearance of tumors after injection, and on day 7 and day 16, tumors were resected with the surrounding skin (when tumors were not visible, a marked site with surrounding tissue collected), fixed in 4% PFA for 48 hours, and processed to paraffin blocks. IHC for GFP, H&E, Ki67, and p-MEK was performed by MSKCC cytology core and reviewed by board certified pathologist.

#### Intracranial injection

50,000 cells were injected at 1 uL/min rate into the brain (injections coordinates: 2 mm right of bregma, 3 mm depth). Mice were randomized (n=3 mice per group) to receive drug treated or DMSO treated cells as a control. Forty-eight hours post-injection, the entire brain was fixed, in 4% PFA for 48 hours. After fixing brain was cut into two halves at the site of injection and processed to paraffin blocks. IHC for GFP, H&E, and p-MEK was performed by MSKCC cytology core and reviewed by board certified pathologist.

All in vivo studies were performed in compliance with institutional guidelines under an approved protocol by the Institutional Animal Care and Use Committee (IACUC). The animals were immediately euthanized as soon as the tumors reached the IACUC set limitations.

### Immunohistochemistry (IHC)

Immunohistochemistry was performed by MSKCC cytology core using Ventana BenchMark ULTRA Automated Stainer. The primary antibodies were hand-applied to the slides followed by an OmniMap HRP multimer detection system (DISCOVERY, Basel, Switzerland). Digital scanning was performed using 3DHISTECH Pannoramic Flash P250” Scanner.

### Bright Field Microscopy

Cells were seeded in 10cm plates in either DMSO or drugs. At 24-hour digital brightfield images were taken using EVOS Brightfield microscope. For Bright field movies cells were plated in 4 well cover glass chamber 24 hour before imaging. Next day glass chamber was mounted in the Zeiss AxioObserver.Z1 stage, treated with DMSO or drug and images recorded for next 24 hours in real time.

### Immunofluorescent and Confocal Microscopy

Cells were seeded in 4 well chamber slides and next day treated with DMSO or drugs. At end points, cells were fixed with 4% PFA, permeabilized by 0.1% Triton X-100 and blocked by 10% goat serum and incubated with primary antibodies overnight and followed by secondary antibodies Cell nuclei were counterstained with DAPI. Images were acquired using Leica TCS SP5” confocal scope. Anti-p-Paxillin (# 2541), anti-Vinculin (# 13901), anti-beta catenin (# 8480) and anti-N-Cadherin (# 13116) antibodies were purchased from cell signaling. Alexa Fluor 488 Phalloidin (#A12379), Goat anti-Rabbit IgG Alexa Fluor 647 (#A 21245), and Goat anti-Rabbit IgG Alexa Fluor 488 (# A32731) antibodies were from Invitrogen.

### RAC1-GTP Assay

GTP-bound RAC1 detected using PAK1-p21-binding domain (PBD) pull-down and Detection Kit (Thermo Scientific, # 16118) as instructed by the manufacturers.

### Phospho-RTK Array

RTK phosphorylation was measured using Proteome Profiler Mouse Phospho-RTK array Kit (R & D Systems, # ARY014) as instructed by the manufacturers.

### Gelatin Invadopodia Assay

Gelatin degradation was measured using QCMTM Gelatin Invadopodia Assay (Green) following manufacturer’s instructions(EMD Millipore, # ECM 670)

### Plasmids and Transfections

PREX1 (# 129619) and PREX2 (# 41555) encoding plasmids were obtained from Addgene. DNA transfections were carried out by using lipofectamine 2000 (Thermo Fisher Scientific, # 11668019 in accordance with the manufacturer’s recommendations.)

### Generation of Stable Cell Lines and Doxycycline inducible lines

The BRAF and RAC1 genes were sub-cloned into TTIGFP-MLUEX vector harboring tet-regulated promoter. Mutations were introduced by using the site-directed Mutagenesis Kit (Stratagene,). HEK293T (RRID: CVCL_0063) cells were used to package virus. Cells were plated in 10 cm tissue culture dish and transfected with 4.5 mg of lentiviral vector (encoding non-targeting, target shRNAs or doxycycline inducible plasmids), 4.5 mg of psPAX2(RRID: Addgene-12260) and 1mg of pVSVG with X-tremeGENE HP (Roche) according to the manufacturer’s protocol. Medium containing recombinant lentiviruses was collected 48 and 72 hours after transfection and filtered through non-pyrogenic filters with a pore size of 0.45 mm (Merck Millipore, Billerica, MA, USA). Samples of these supernatants were applied immediately to target cells together with Polybrene (Sigma-Aldrich, St. Louis, MO, USA) at a final concentration of 6 mg/ml, and supernatants were incubated with cells for 24 hr. After infection, cells were placed in fresh growth medium and cultured as usual. Selection with 2.5 ug/ml puromycin was initiated 48 h after infection for 3 days. Hairpins were purchased from Sigma Aldrich and are listed below: Short hairpins used against RAC1 are (#TRCN0000055191), and (#TRCN00000301589). Short hairpins used against PTEN are (#TRCN0000028992) and (# TRCN00000322486). The control hairpin used is (# SH016)

## STATISTICAL ANALYSIS

The details of statistical analysis of experiments can be found in the figure legends. Statistical analysis of differences between samples was performed using two-tailed Student’s t-tests, and p < 0.05 was defined as significant. One-way ANOVA analysis was performed to compare the means of more than two groups. All analysis was conducted using GraphPad Prism (RRID: SCR_002798). Independent experiments were conducted with a minimum of two biological replicates per condition to allow for statistical comparison.

Data is shown as mean ± SEM.

---

**Figure 7:**
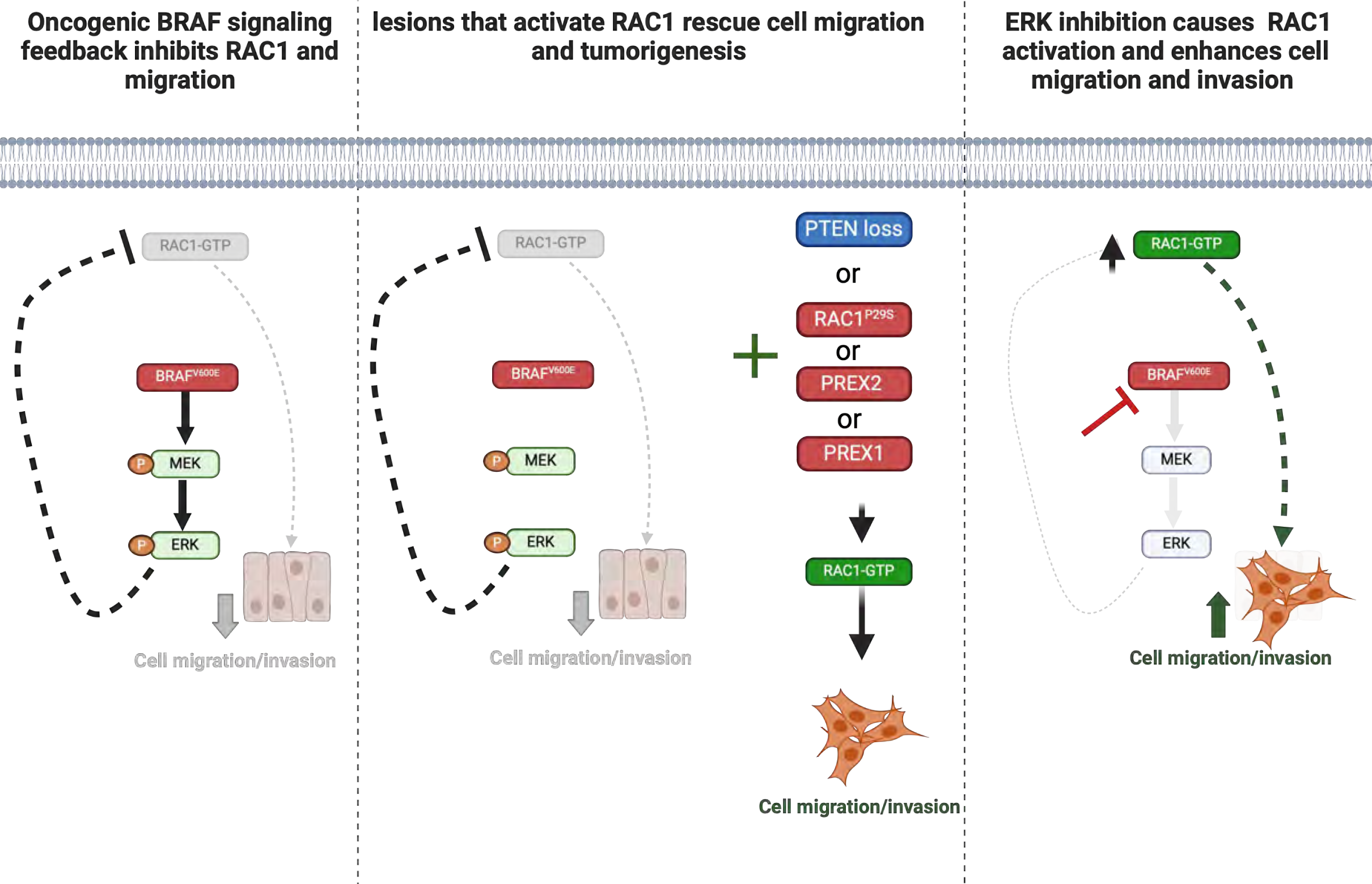
Grapical abstract depicting how oncogenic BRAF^V600E^ shapes tumorigenesis. Oncoprotein BRAF^V600E^ feedback inhibits cell migration and invasion, processes necessary for malignant transformation(1st panel). Secondary genetic lesions rescue migration, some by activating RAC1, and support tumorigenesis (2rd panel). Inhibition of ERK signaling relieves this feedback, and induces RAC1 activation and cell migration(3rd panel).

## Notes

### Competing Interest Statement

The authors have declared no competing interest.

### Summary of Updates

Title, Discussion section and graphical Abstract has been revised

