## Supplementary material for "Tumorigenesis driven by the BRAF^V600E^ oncoprotein requires secondary mutations that overcome its feedback inhibition of migration and invasion": supplemental data.pdf

Supplemental Figure1: BRAF$^{V600E}$ -driven ERK signaling inhibits cell migration in vitro.

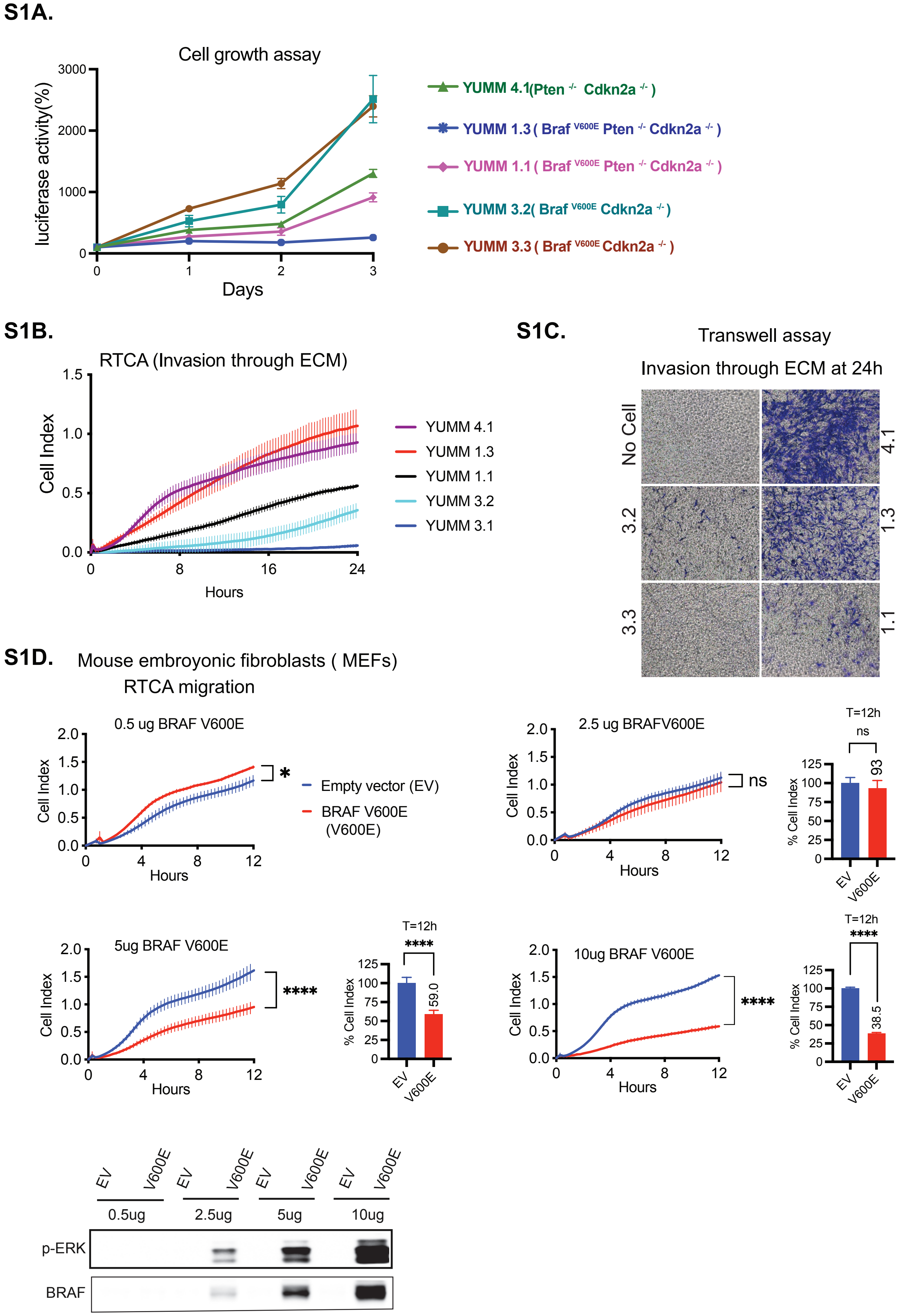

Supplemental Figure1: BRAF$^{V600E}$-driven ERK signaling inhibits cell migration in vitro.

**S1E**

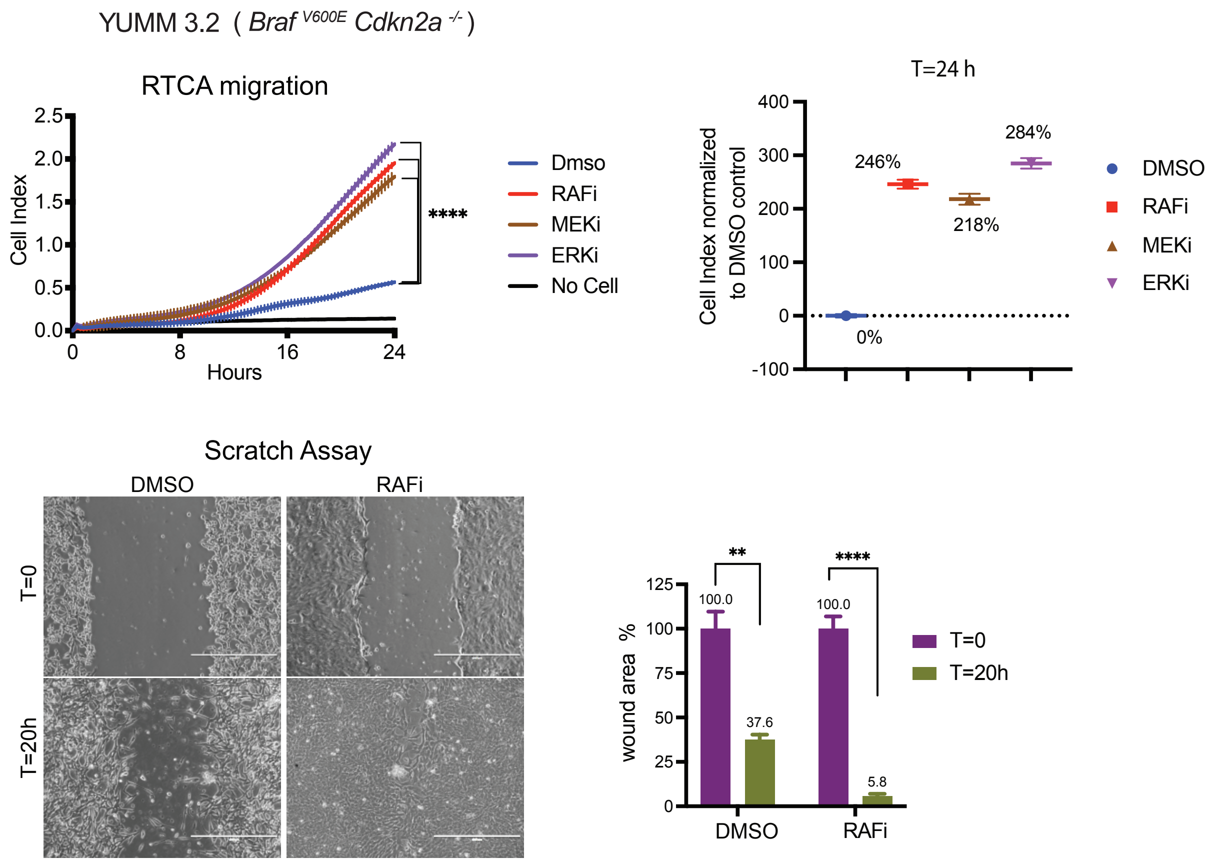


**S1F.**

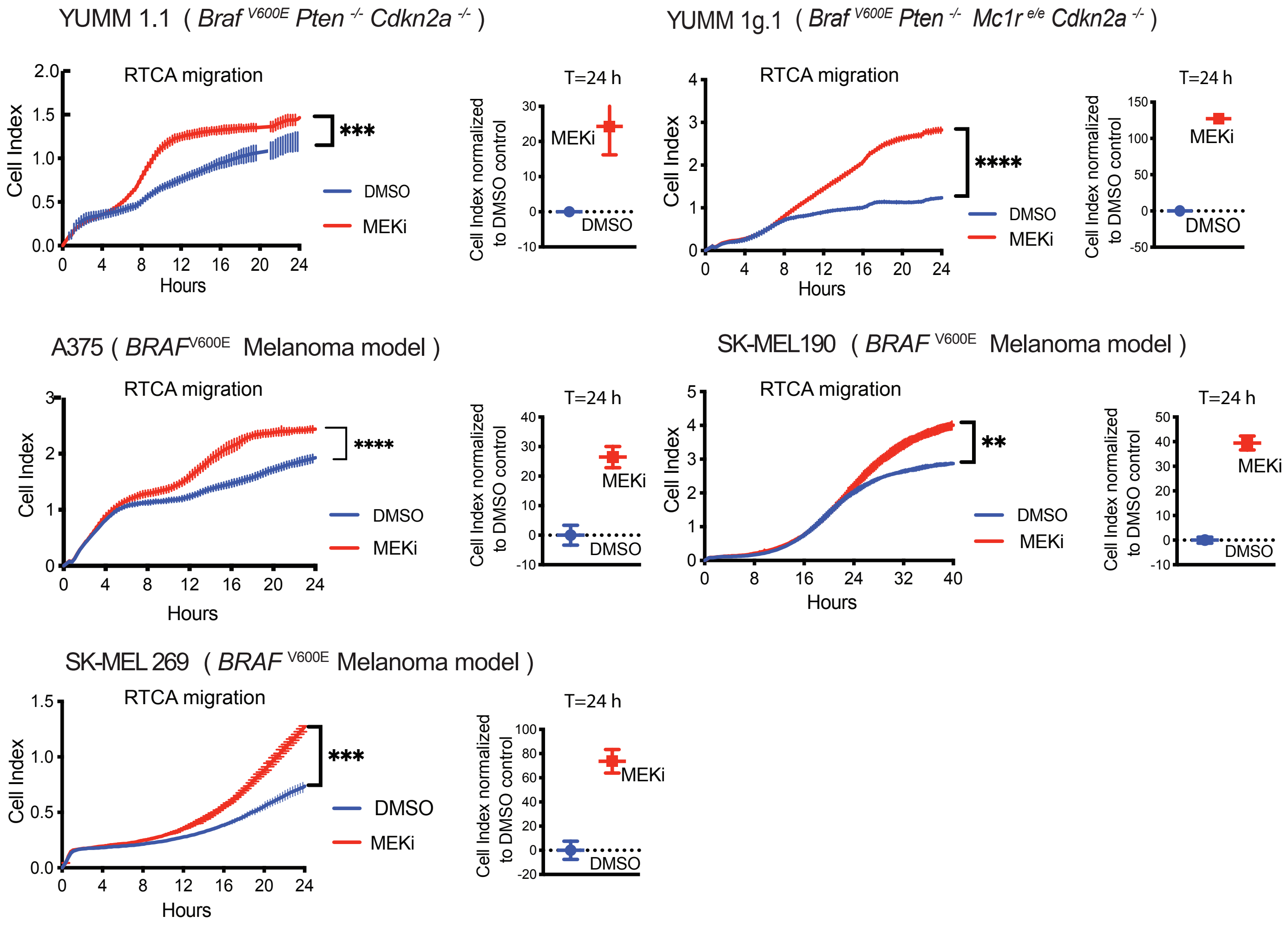

Supplemental Figure1: BRAF$^{V600E}$ -driven ERK signaling inhibits cell migration in vitro.

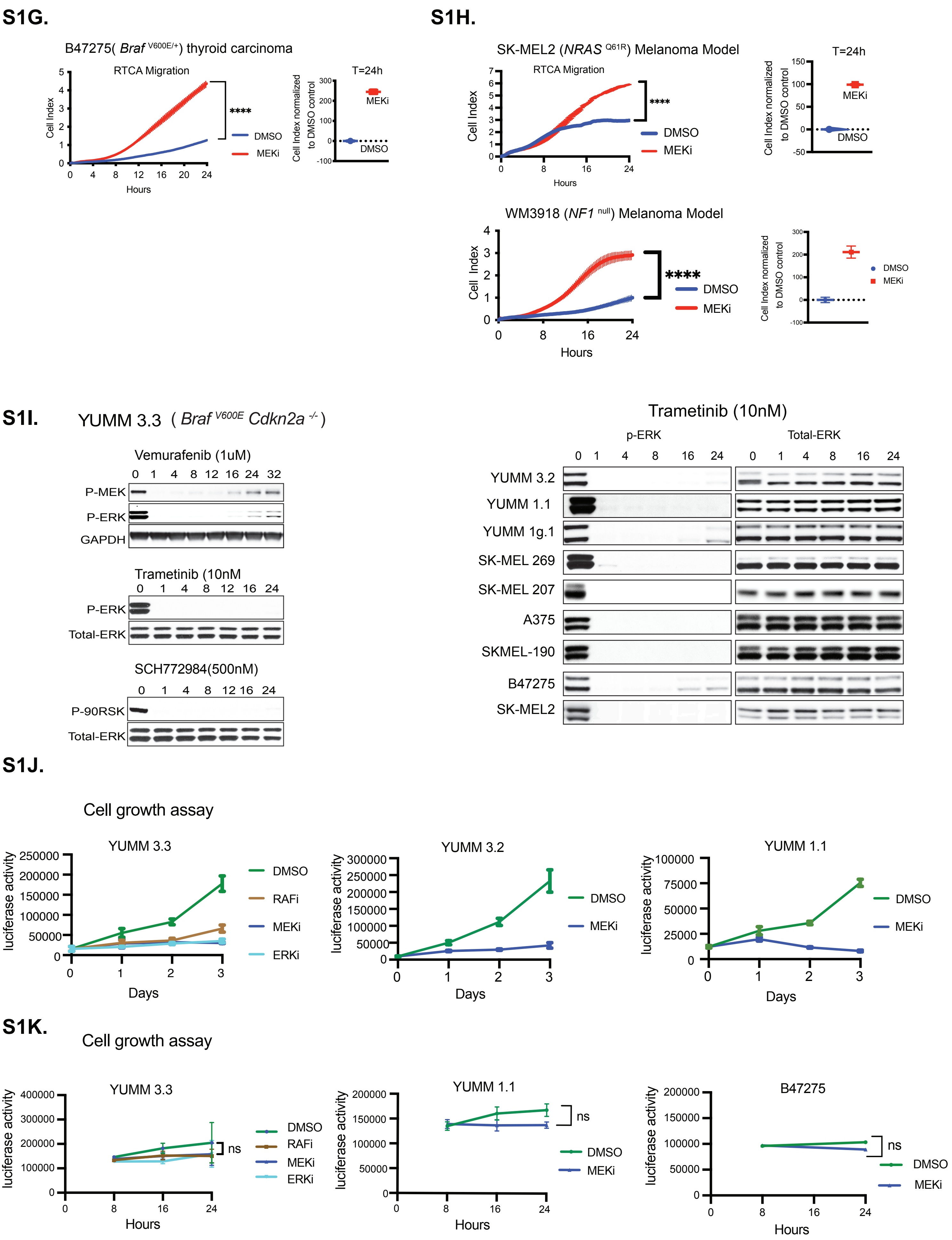

**Figure SF1, related to Figure 1.**
**Supplemental Figure 1: BRAF$^{V600E}$-driven ERK signaling inhibits cell migration in vitro.**

**S1A.** YUMM lines YUMM 4.1, YUMM 1.3, YUMM 1.1, YUMM 3.3, and YUMM 3.2 were seeded in 96 well plates at 1000 cells/well. Cell growth was measured at indicated times using CellTiter-Glo Cell Viability Assay. (n=6, Error bars represent mean ± SE).

**S1B.** Invasion curves of YUMM 4.1, YUMM 1.3, YUMM 1.1, YUMM 3.3, and YUMM 3.2 were generated by seeding the cells in ECM-coated upper chambers. The curves with vertical error bars represent mean ± SEM (n=3)

**S1C.** For transwell assay, cells were seeded in ECM-coated upper chambers, and the underbelly of the upper chambers was fixed at the indicated times and visualized by Crystal Violet (CV). Representative images from 3 independent experiments are shown.

**S1D.** MEFs were transfected with an empty vector or BRAF$^{V600E}$ expressing Vector with indicated plasmid concentrations. Thirty-six hours after transfection, cells were seeded to generate migration curves. The curves with vertical error bars represent mean ± SEM (n=3). The bar graph depicts the percent change in cell index at 12 hours compared to the empty vector. WCL was analyzed by IB for indicated proteins.

**S1E.** Migration curves of YUMM 3.2 treated with either DMSO, 1uM Vemurafenib (RAFi), 10nM Trametinib (MEKi), or 500nM SCH 772984 (ERKi) treatment. p values are based on ordinary one-way ANOVA analysis. Cell index at 24 hours was normalized to DMSO control. Migration curves with vertical error bars represent mean ± SEM (n=3). For Scratch Assay, YUMM 3.2 cells were plated with DMSO or 1uM Vemurafenib. Twenty-four hours after plating, a scratch was made. Bright-field image of the scratch was taken at T=0h (within 30min of wounding) and T=20h (20 hours after wounding). The bar graph depicts the open wound area. Here, we showed a representative image of 3 independent experiments.

**S1F.** Migration curves of YUMM 1.1, YUMM 1g.1, A375, SK-MEL190, and SK-MEL269 cells were generated upon treatment with either DMSO or 10nM Trametinib. Migration curves with vertical error bars represent mean ± SEM (n=3). The cell index at 24 hours was normalized to DMSO control.

**S1G.** Migration curves of *Braf*$^{V600E}$ thyroid carcinoma mouse cell line B47275 were generated upon treatment with either DMSO or 10nM Trametinib. Migration curves with vertical error bars represent mean ± SEM (n=3). Cell index at 24 hours was normalized to DMSO control.

**S1H**. Migration curves of NRAS mutant melanoma cell model SK-MEL2 and NF1 null melanoma model WM3918 were generated upon treatment with either DMSO or 10nM Trametinib. Migration curves with vertical error bars represent mean ± SEM (n=2). Cell index at 24 hours was normalized to DMSO control.

**S1I.** YUMM 3.3 cells were treated with either DMSO, 1uM Vemurafenib, 10nm Trametinib, or 500nM SCH 772984 for indicated times, and WCL was analyzed by IB for indicated proteins. The mentioned cell lines were treated with 10nM Trametinib for indicated times and analyzed by IB.

**S1J**. YUMM 3.3, 3.2, and 1.1 cells were seeded in 96 well plates at 1000 cells/well. Cell growth was measured at indicated times using CellTiter-Glo Cell Viability Assay. Error bars represent mean ± SE (n=6).

**S1K**. For the cell proliferation assay, cells were seeded in 96 well plates and treated with either DMSO, 1uM Vemurafenib,10nM Trametinib, or 500nM SCH 772984 at the time of plating. Cell growth was measured at indicated times using CellTiter-Glo Cell Viability Assay. Error bars represent mean ± SE (n=3). Migration curves with vertical error bars represent mean ± SEM (n=3). p values are based on a two-tailed unpaired Student's t-test.

*P*<0.0001 is shown as ****, P<0.001 as ***, P< 0.01 as **, and *P*>0.05 as ns.

Supplemental figure 2: BRAF$^{V600E}$- driven ERK signaling inhibits cell migration and invasion in vivo.

**A**

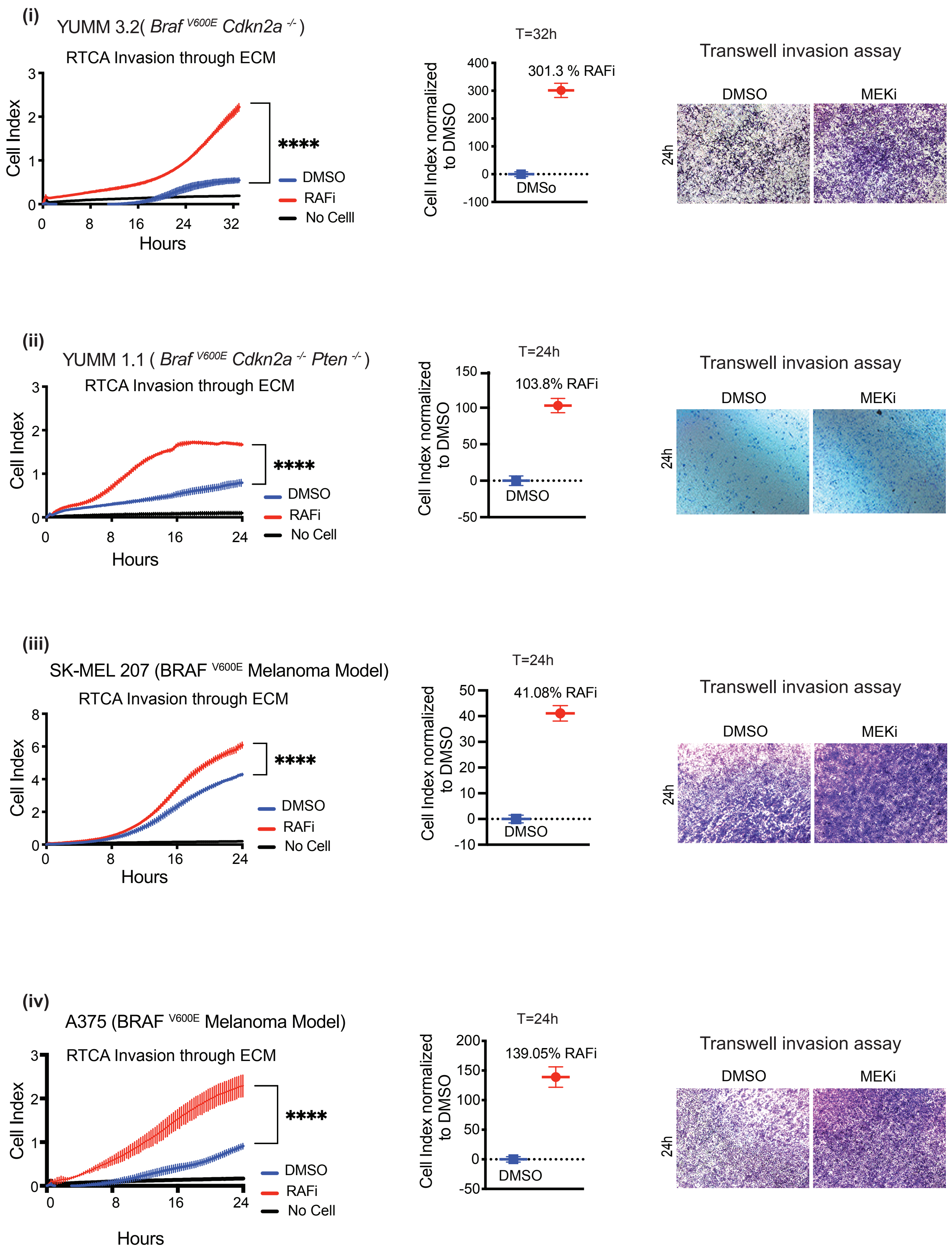

Supplemental figure 2: $BRAF^{V600E}$- driven ERK signaling inhibits cell migration and invasion in vivo.

**B**

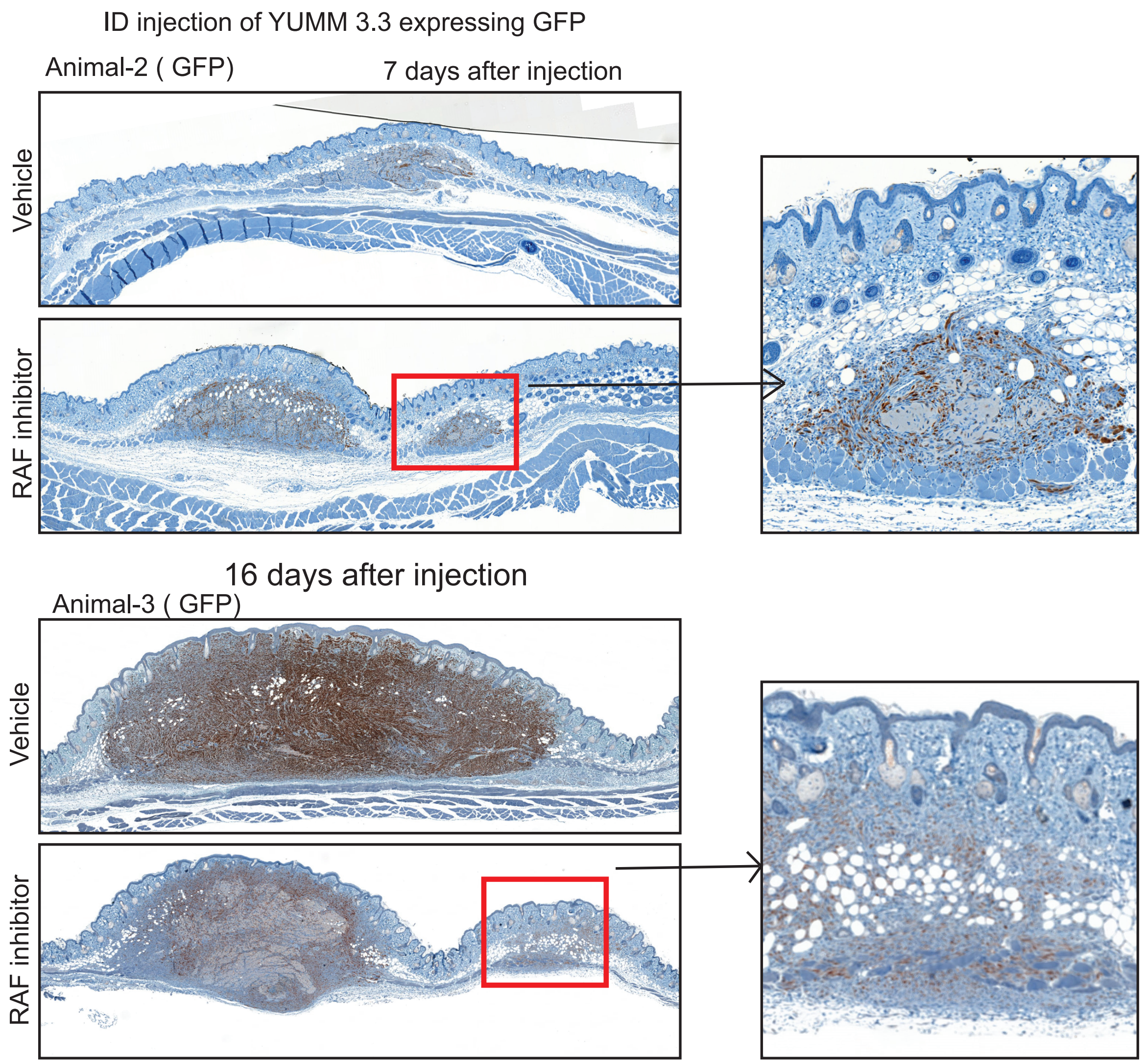


**C**

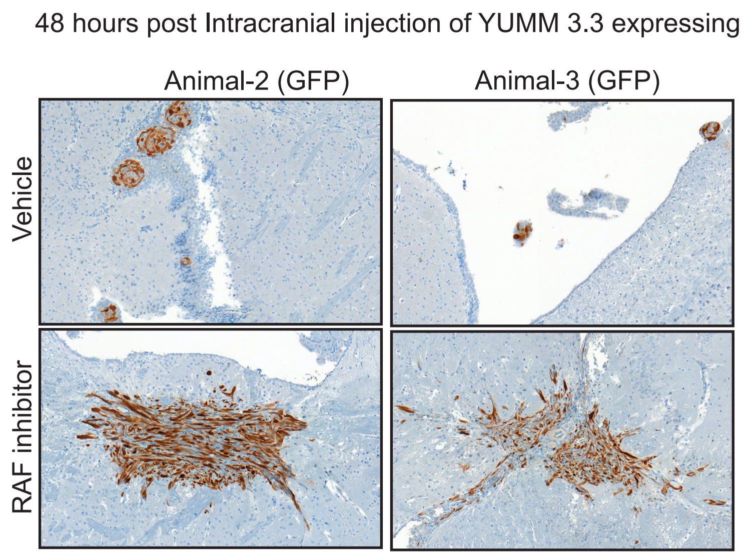

**Figure SF2, related to Figure 2.**
**Supplemental Figure 2: BRAF$^{V600E}$-driven ERK signaling inhibits migration and invasion in vivo.**

**S2A**. Invasion traces of YUMM 3.2, YUMM 1.1, SK-MEL207, and A375 were generated using ECM-coated plates with either DMSO or 1uM Vemurafenib. The invasion cell index at 24h was normalized to the DMSO control. For transwell assay ECM-coated seeding plates were used, and cell invasion was assessed at indicated times, followed by CV staining.

**S2B.** YUMM 3.3 cells were pre-treated with either DMSO or 1uM PLX4720 (RAF inhibitor) for 24 hours and injected intradermally. Mice were treated with either DMSO or 7.5mg/kg PLX4720 24-hour post-injection. 7- & 16-day post-injection tumors were collected, and histopathological sections were stained with GFP. The red box shows secondary foci evocative of microsatellitosis. The inset displays tumor microsatellitosis at higher magnification.

**S2C**.GFP expressing Yumm3.3 cells were pre-treated with either DMSO or 1uM Vemurafenib for 24 hours and injected intracranially. 48 hours post-injection brain was collected, and histological sections were stained with GFP. While control tumors remained confined within the ventricular cavity, RAFi-treated melanomas crossed the ependymal lining and invaded the brain parenchyma as irregular nodules with jagged tumor borders and a single cell pattern of growth.

Suplemental Figure 3: ERK pathway inhibition causes mesenchymal transition.

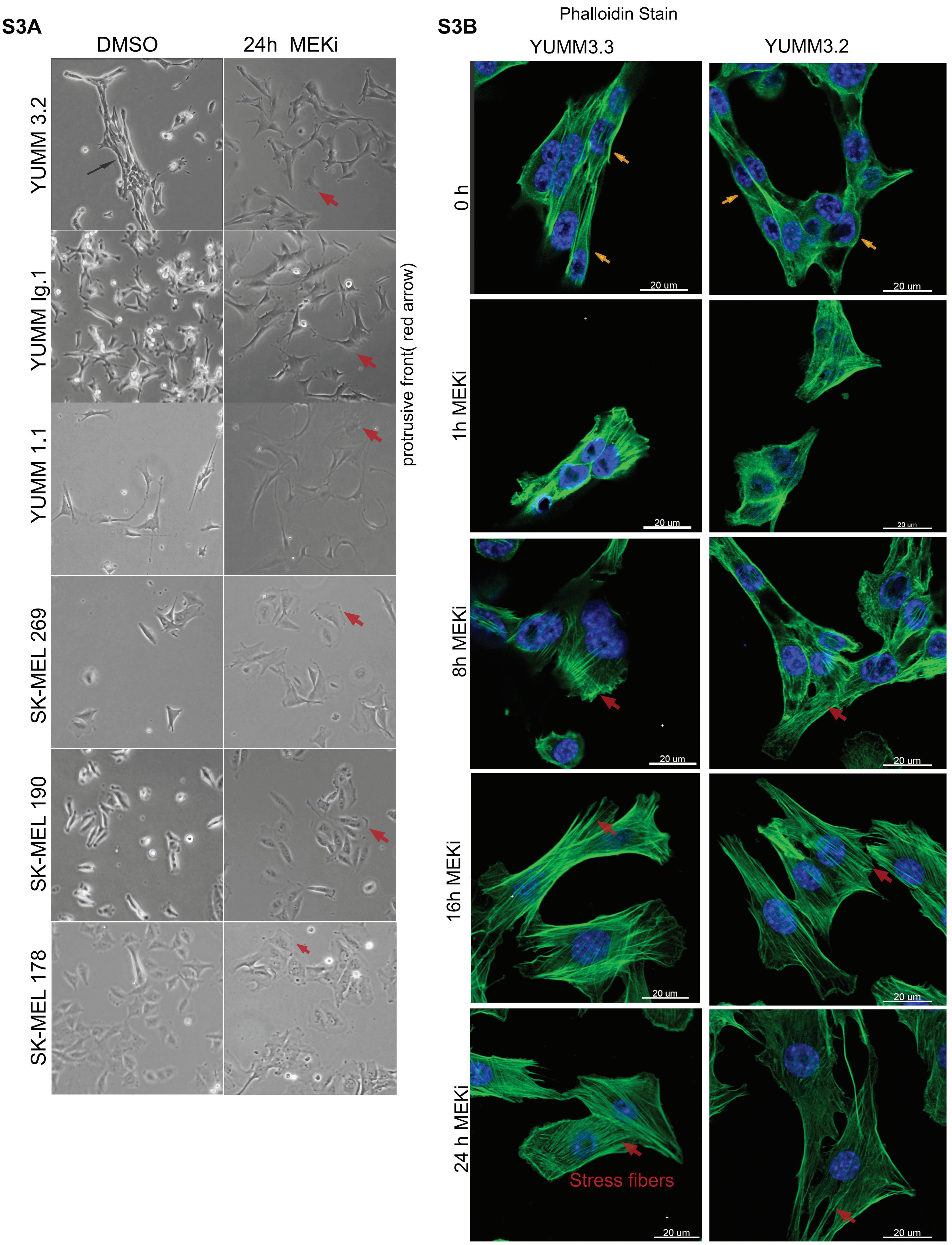

Suplemental Figure 3:  ERK pathway inhibition causes mesenchymal transition.

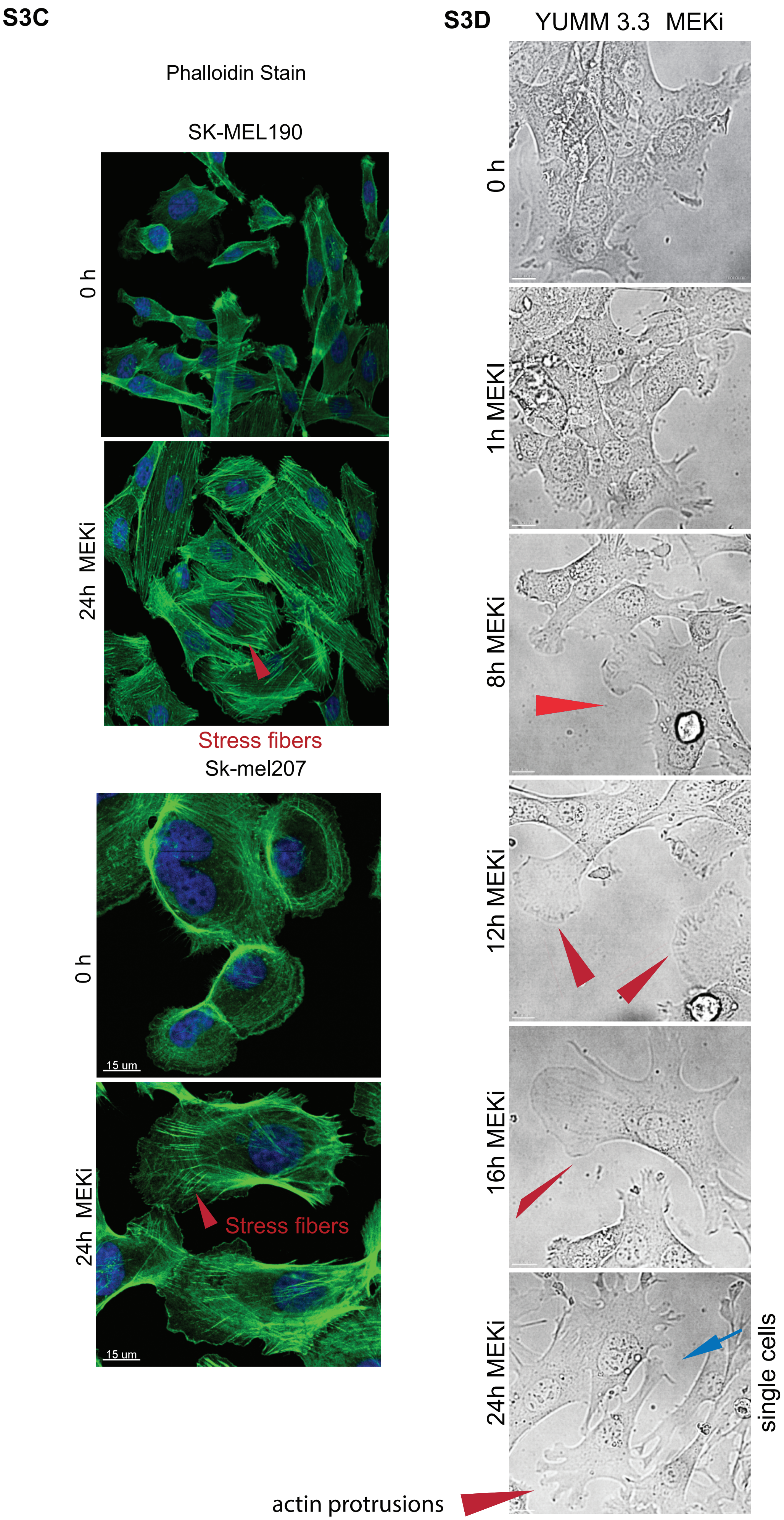

**Figure SF3, related to Figure 3.**
**Supplemental Figure 3: ERK pathway inhibition triggers mesenchymal transition.**

**S3A**.Representative bright-field images of indicated cell lines 24 hours after their exposure to either DMSO or 10nM Trametinib. The red arrows show actin protrusions at the leading edge. Magnification 10X

**S3B**.Representative confocal images of YUMM 3.3 and YUMM 3.2 after exposure to DMSO or 10nM Trametinib. Cells were fixed at the indicated time after treatment, and actin filaments were visualized with a phalloidin stain (green). The yellow arrow shows the cortical actin. The red arrow shows the induction of stress fibers. Nuclei were stained with DAPI. Scale bar, 20uM.

**S3C**. Representative confocal images of SK-MEL 190 and SK-MEL207 treated with either DMSO or 10nM Trametinib (24h). Actin filaments were visualized with a phalloidin stain (green). The red arrow shows the induction of stress fibers. Nuclei were stained with DAPI. Scale bar, 15uM.

**S3D**.Brightfield images of YUMM 3.3 cells were taken at the indicated time upon treatment with 10nM Trametinib. The red arrow shows a gradual sequence of lamellipodia formation, and the blue arrow shows cell dispersion. Magnification 10X

**S video1a:** Confocal Z-stack image of YUMM3.3 treated with DMSO.

**S video1b:** Confocal Z-stack image of YUMM3.3 treated with 10nM Trametinib for 24 hours.

**S Video-2:** Time-lapse video of SK-MEL 207 with DMSO or 10nM Trametinib. The red arrow follows the control cell, while the magenta and blue arrow follow the MEK inhibitor-treated cells.

Supplemental Figure 4: RAC1 activation causes mesenchymal transition and enhances cell motility upon ERK inhibition.

**S4A.**

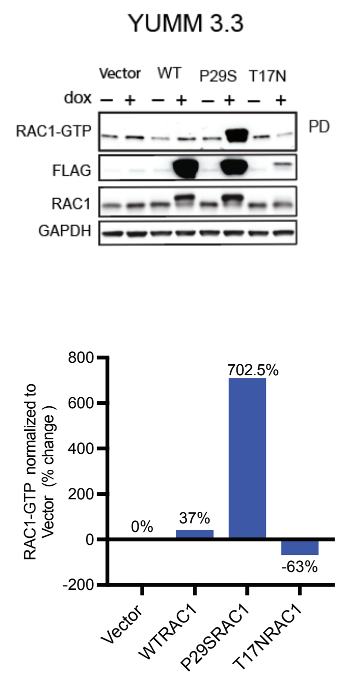


**S4B.**

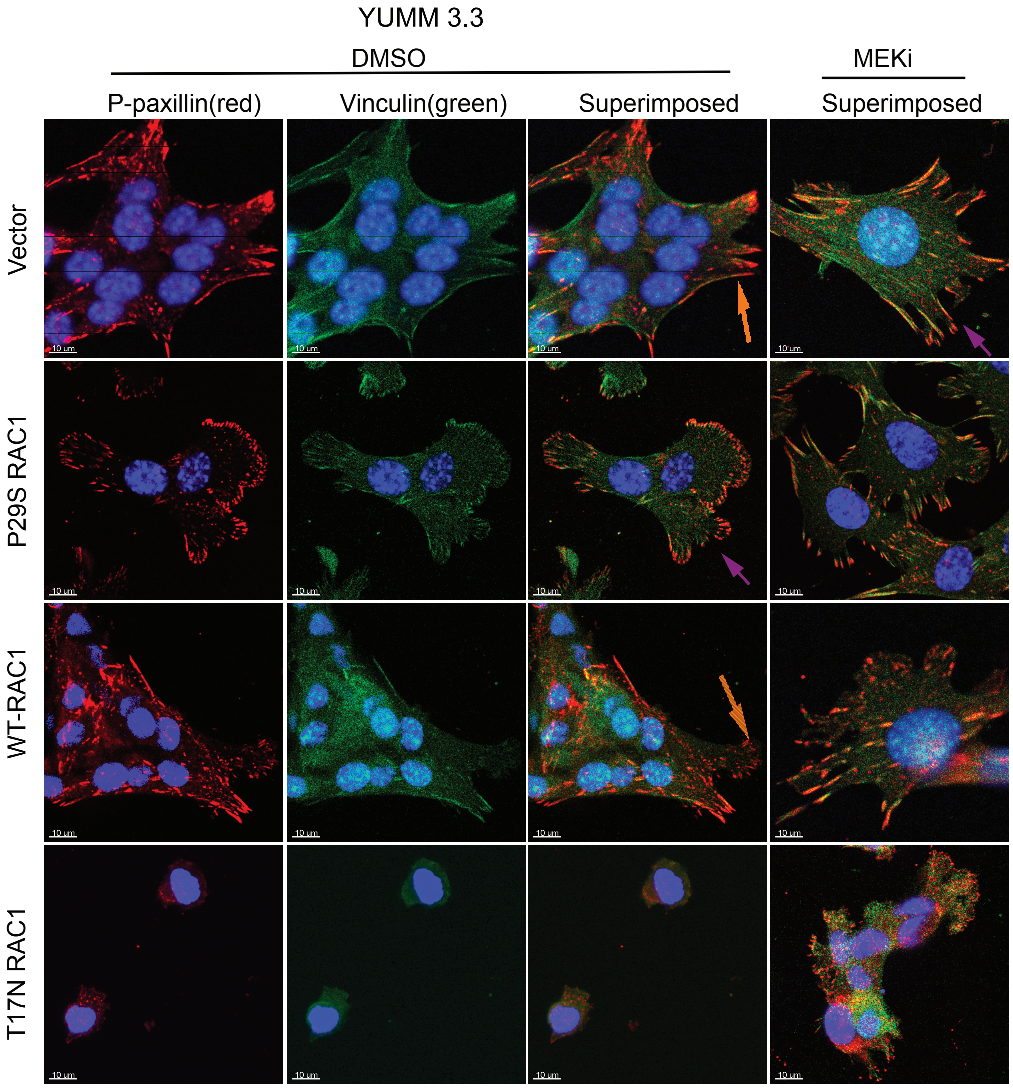


**S4C.**

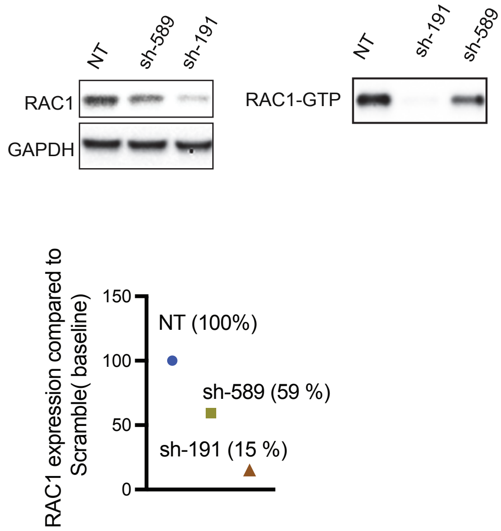


**S4D.**

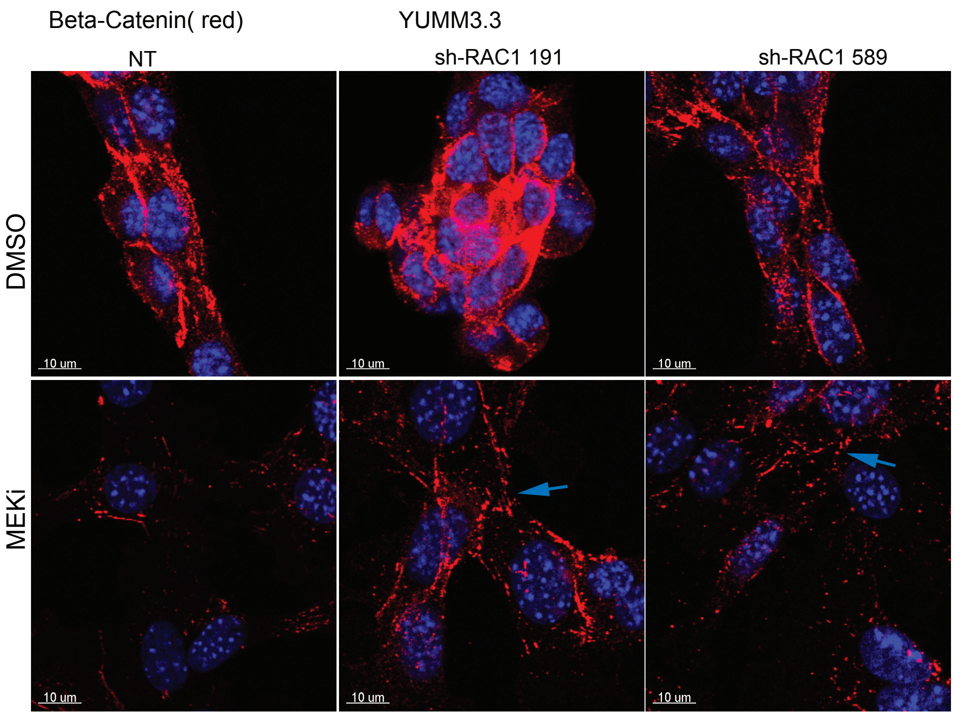

Supplemental Figure 4: RAC1 activation causes mesenchymal transition and enhances cell motility upon ERK inhibition.

S4E.

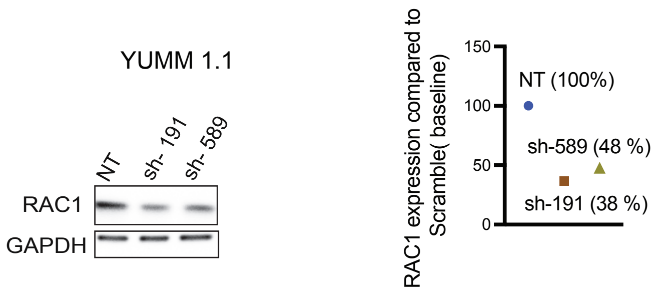


S4F.

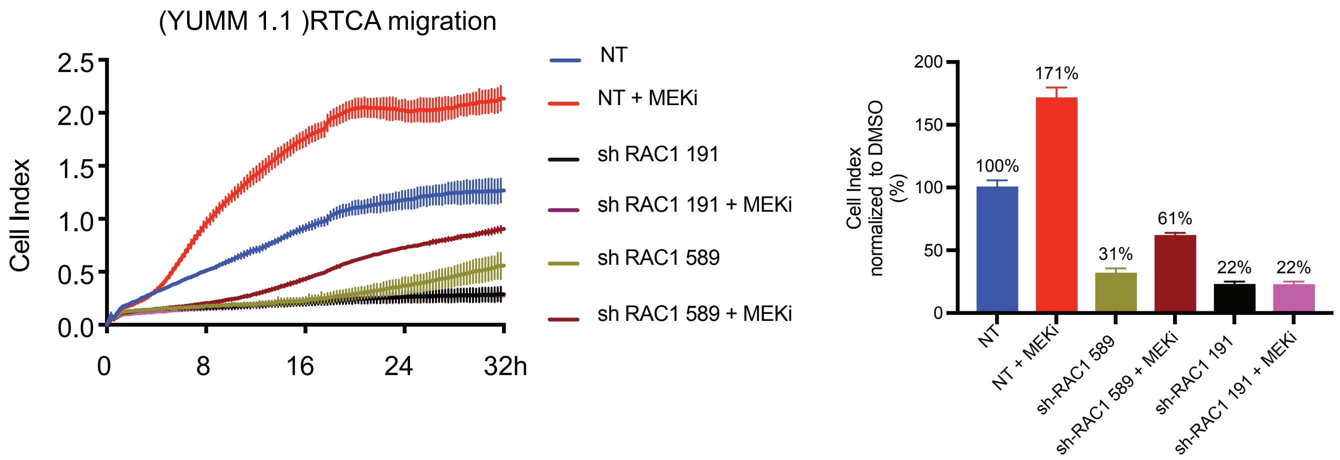


S4G.

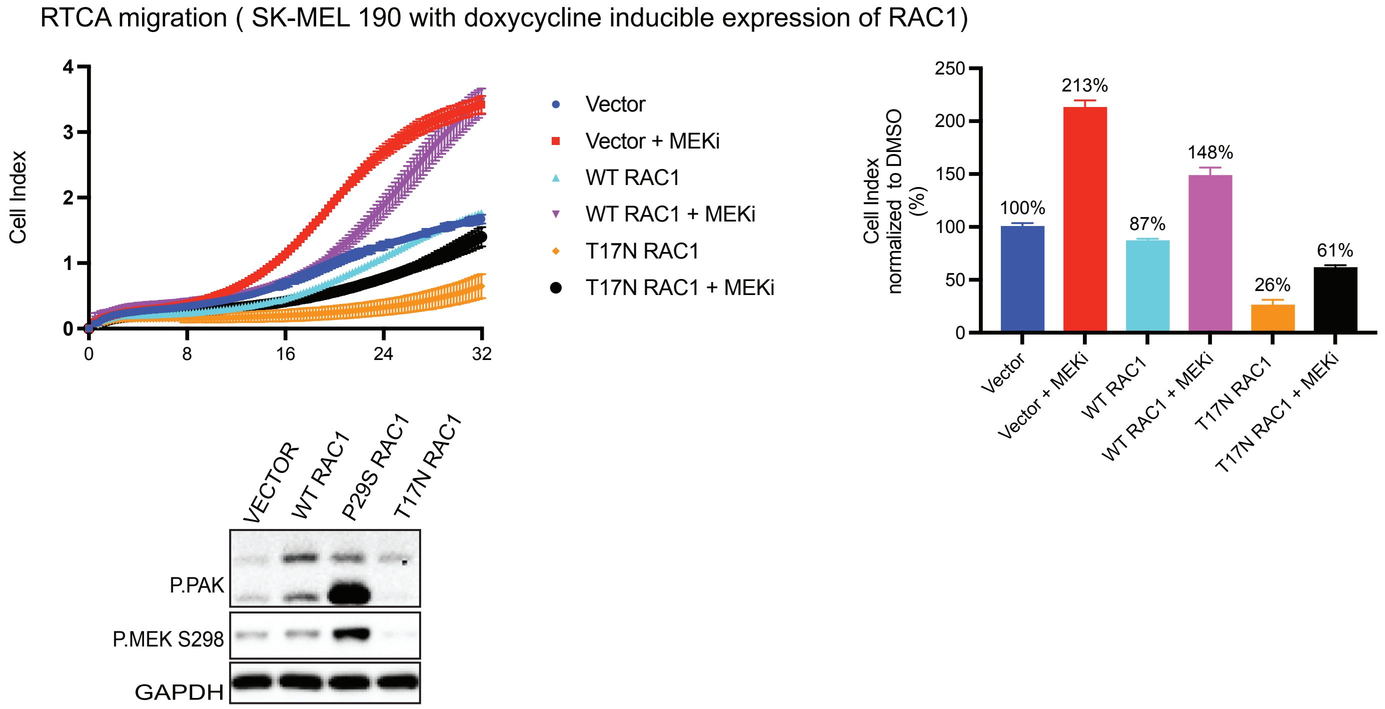

**Figure SF4, related to Figure 4.**
**Supplemental Figure 4: RAC1 activation causes mesenchymal transition and enhances cell motility upon ERK inhibition**

**S4A**.YUMM 3.3 with inducible expression of empty vector, WT RAC1, or constitutively active mutant P29S RAC1 (1ug/ml doxycycline 24h) were treated with DMSO or 10nM Trametinib. Twenty-four hours after treatment, WCL was collected and assessed for changes in RAC1.GTP and pERK as described in Fig 4A legend. Quantification of RAC1 GTP was normalized to a DMSO-treated empty vector.

**S4B.**YUMM 3. with inducible expression of empty vector, WT RAC1, $RAC1^{P29S,}$ or the dominant negative $RAC1^{T17N}$ (1ug/ml doxycycline 24h) were treated with DMSO or 10nM Trametinib for 24 hours. Cells were fixed, and the focal adhesion complex was visualized with vinculin (green) or phosphorylated paxillin Y118 (red). The superimposed image (orange) shows the co-localization of vinculin and phosphorylated paxillin. Scale bar 10uM

**S4C.**Downregulation of RAC1 expression in YUM3.3 cells expressing short hairpins against RAC1. WCL were collected from YUMM 3.3 cells expressing either NT or one of two different short hairpins against RAC1, sh-RAC1 191, and sh-RAC1 589 and assayed for RAC1 expression, GAPDH and activated RAC1 as in 4A. Quantification of RAC1 expression was normalized to YUMM 3.3 expressing NT hairpin.

**S4D**.Visualization of intercellular junction with a beta-catenin stain. YUMM 3.3 cells expressing NT or short hairpins against RAC1 (sh-RAC1 191 and sh-RAC1 589) were treated with DMSO or 10nM Trametinib. Upon 24 hours of treatment, cells were fixed, and intercellular junctions were visualized with a beta-catenin stain (red). Scale bar 10uM

**S4E.** WCLs of YUMM 1.1 cells expressing either NT or short hairpins against RAC1 (sh-RAC1 191 and sh-RAC1 589) were collected, and RAC1 expression was determined by IB with indicated antibodies. Quantification of RAC1 expression was normalized to YUMM 1.1 expressing NT.

**S4F**.RAC1 expression is required for the migration of a $BRAF^{V600E}PTEN^{-/-}$ cell and for its response to MEK inhibition. Migration of YUMM 1.1 cells expressing NT or short hairpins against RAC1 (sh-RAC1 191 and sh-RAC1 589) was determined with or without 10nM Trametinib. Vertical error bars on the cures represent mean $\pm$ SEM (n=3). The bar graph shows the percent cell index normalized to DMSO control at 24 hours.

**S4G**. Migration curves of SK-MEL 190 with inducible expression of empty vector, WT RAC1, or $RAC1^{T17N}$ (1ug/ml doxycycline 24h) with or without 10nM

Trametinib. Curve with vertical error bars represents mean ± SEM (n=3). The Bar graph shows percent cell index is normalized to DMSO control at 24 hours. WCL of the indicated cells was collected 24h after 1ug/ml doxycycline and immunoblotted with antibodies for the indicated protein.

Supplemental Figure 5: Relief of ERK-dependent feedback potentiates RTK signaling, which in turn activates RAC1 and cell migration.

**S5A**

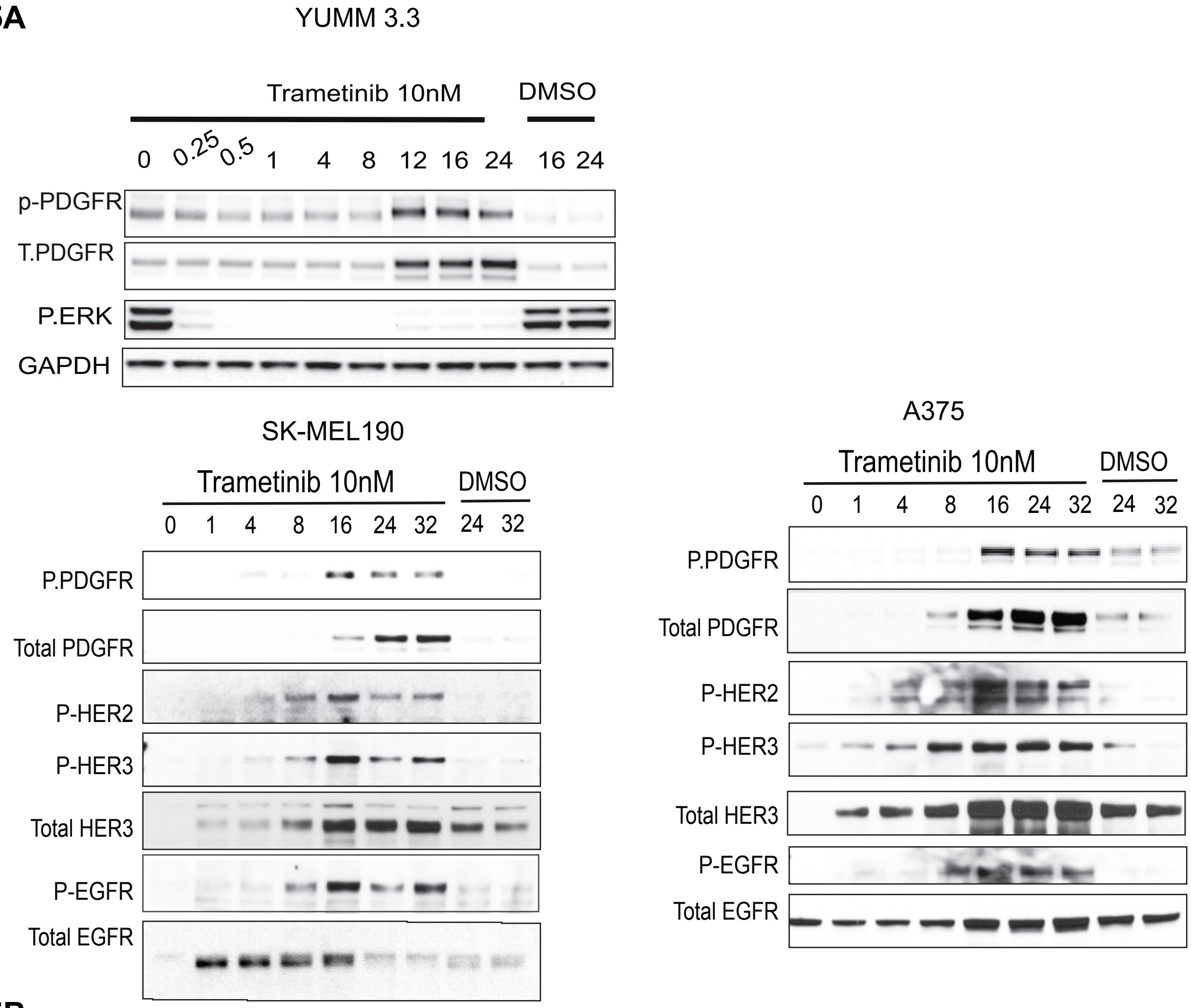


**S5B**

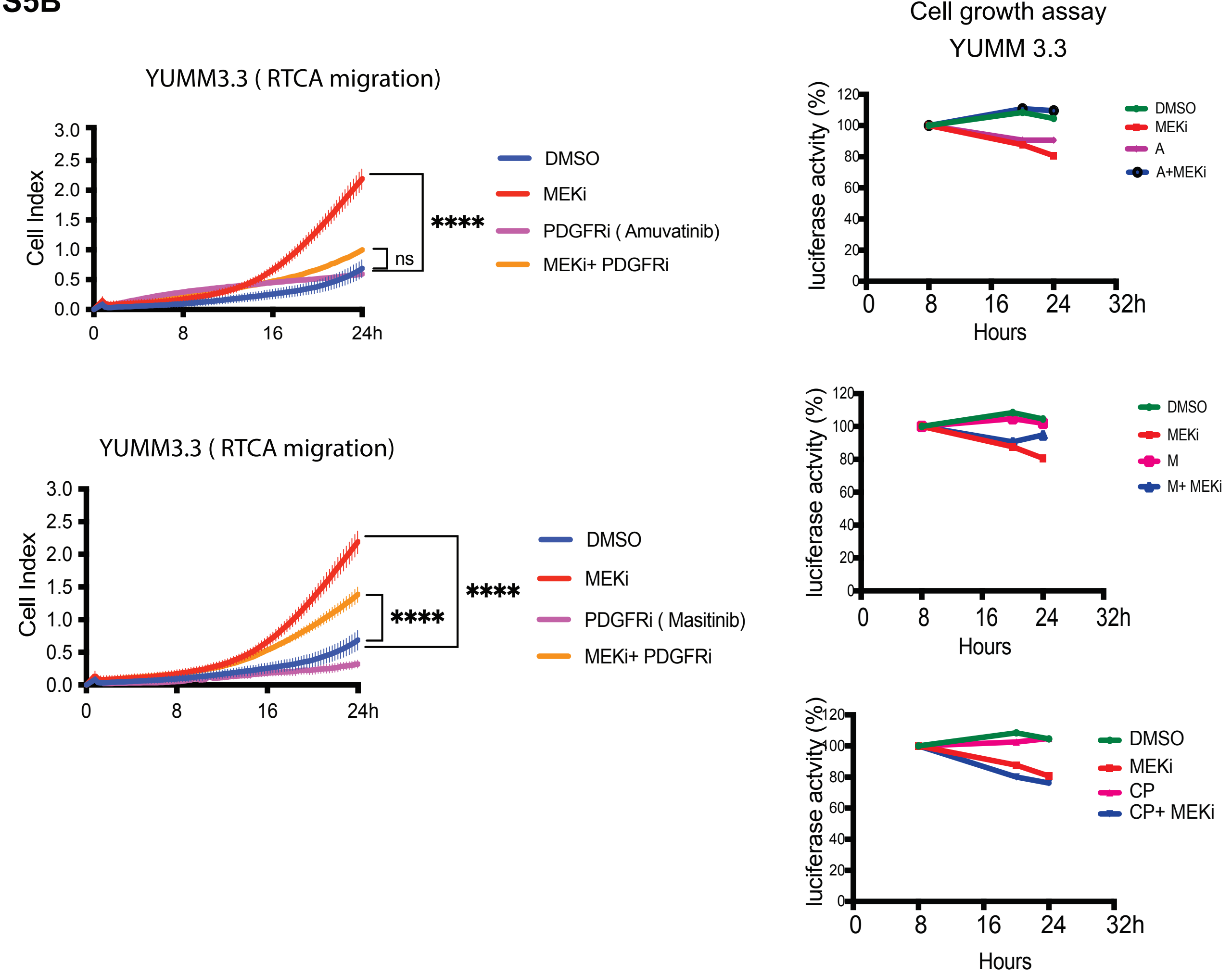

**Figure SF5, related to Figure 5.**
**Supplemental Figure 5: Relief of ERK-dependent feedback potentiates RTK signaling, which in turn activates RAC1 and cell migration**

**S5A.** YUMM 3.3, A375, and SK-MEL190 cells were treated with either DMSO or 10nM Trametinib for the indicated time, and WCLs were IB for indicated proteins.

**S5B.** Migration of YUMM 3.3 cells treated with either DMSO, 10 nM Trametinib as single agent and 5uM Amuvatinib (A), or 5uM Masitinib (M) as single agents or in combination with 10nmM Trametinib. Cell growth was measured at indicated times using CellTiter-Glo Cell Viability Assay. (n=3, Error bars represent mean ± SE).

Supplemental Figure 6 :BRAF$^{V600E}$ melanoma are enriched with lesions that rescue cell motility.

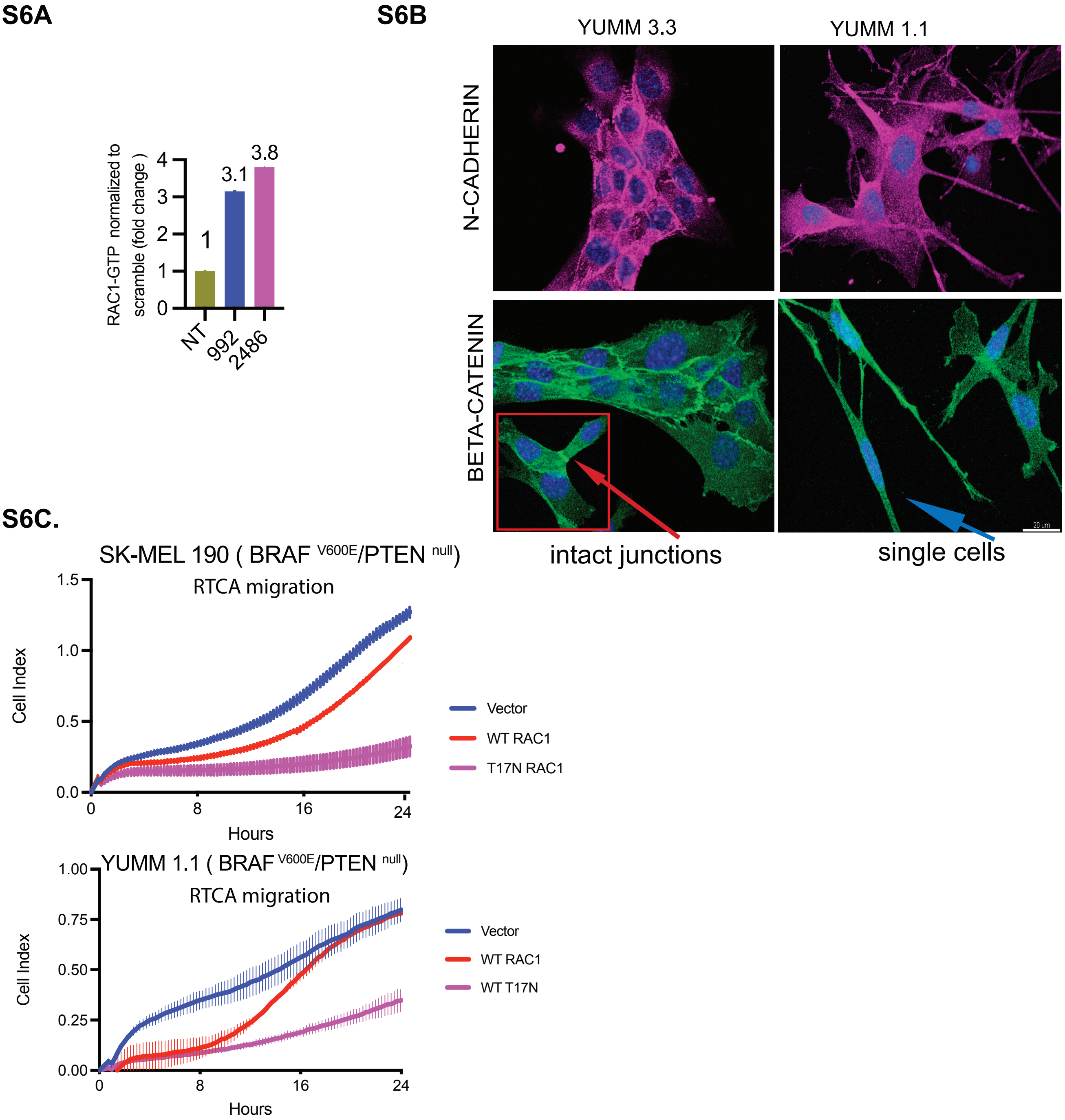

**Figure SF6, related to Figure 6.**
**Supplemental Figure 6: BRAF$^{V600E}$ melanoma is enriched with lesions that rescue cell motility.**

**S6A.**RAC1-GTP levels of the indicated cells were quantified and normalized to YUMM 3.3, expressing NT.

**S6B**.YUMM 3.3 and YUMM 1.1 cells were fixed, and intercellular junctions were visualized with either β-catenin(green) or N-cadherin (magenta) stains. The arrow shows intact or absence of junctions. The inset shows an intact junction in YUMM 3.3. Scale bar 20uM

**S6C**.Migration curves SK-MEL 190 and YUMM 1.1 expressing wildtype RAC1 or dominant negative mutant RAC1$^{T17N}$ (1ug/ml doxycycline 24h). Curve with vertical error bars represents mean ± SEM (n=3).
